# Repeated evolution of circadian clock dysregulation in cavefish populations

**DOI:** 10.1101/2020.01.14.906628

**Authors:** Katya L. Mack, James B. Jaggard, Jenna L. Persons, Courtney N. Passow, Bethany A. Stanhope, Estephany Ferrufino, Dai Tsuchiya, Sarah E. Smith, Brian D. Slaughter, Johanna Kowalko, Nicolas Rohner, Alex C. Keene, Suzanne E. McGaugh

**Affiliations:** Biology, Stanford University, Stanford, CA, USA; Department of Biological Sciences, Florida Atlantic University, Jupiter, FL, USA; Stowers Institute for Medical Research, Kansas City, MO, USA; Ecology, Evolution, and Behavior, University of Minnesota, Saint Paul, MN, USA; Wilkes Honors College, Florida Atlantic University, Jupiter FL, USA; Department of Molecular and Integrative Physiology, The University of Kansas Medical Center, Kansas City, KS, USA

## Abstract

Circadian rhythms are nearly ubiquitous throughout nature, suggesting they are critical for survival in diverse environments. Organisms inhabiting environments with arrhythmic days, such as caves, offer a unique opportunity to study the evolution of circadian rhythms in response to changing ecological pressures. Here we demonstrate that the cave environment has led to the repeated disruption of the biological clock across multiple populations of Mexican cavefish, with the circadian transcriptome showing widespread reductions in rhythmicity and changes to the timing of the activation/repression of genes in the core pacemaker. Then, we investigate the function of two genes with decreased rhythmic expression in cavefish. Mutants of these genes phenocopy reductions in sleep seen in multiple cave populations, suggesting a link between circadian dysregulation and sleep reduction. Altogether, our results reveal that evolution in an arrhythmic environment has resulted in dysregulation to the biological clock across multiple populations by diverse molecular mechanisms.

## Introduction

Circadian rhythms that maintain 24-hour oscillations in physiology and behavior are nearly ubiquitous in nature^1,2^. Considered an adaptive mechanism for organisms to anticipate predictable changes in their environment^3,4^, the biological clock coordinates diverse biological processes, from the sleep-wake cycle and metabolism in animals to growth and photosynthesis in plants^5–7^. In the majority of organisms, the biological clock is endogenous, but environmental time-cues (“zeitgebers”) including light and temperature play a key role in synchronizing the clock with an organism’s external environment^8–11^. Subjecting animals to light-dark cycles that differ from that of a 24-hour day has profound impacts on organismal health, including reduced performance, increased illness, and decreased longevity^12–14^. Consequently, systems where these environmental zeitgebers have been lost provide a unique opportunity to examine how ecology drives evolution in the biological clock, and characterize how clock evolution affects downstream physiology and behavior^15^. Here, we examine daily expression changes across the transcriptomes of cavefish populations that evolved under darkness and relatively low daily temperature variation^16,17^.

The Mexican tetra, *Astyanax mexicanus*, exists as surface populations that live in rivers with robust light and temperature rhythms, and at least 30 cave populations that live in perpetual darkness with limited fluctuations in temperature or other environmental cues^18^. The existence of multiple cave populations of independent origin and conspecific surface populations provides a powerful comparative framework to study phenotypic evolution. Cave populations of *A. mexicanus* have repeatedly evolved a suite of traits related to a cave-dwelling lifestyle, including degenerate eyes^19–21^, reduced pigmentation^22–25^, and changes in metabolism and behavior^26–35^.

Circadian rhythms and sleep behavior are also substantially altered in cavefish^27,36–38^. While cavefish largely maintain locomotor and physiological rhythms in light-dark conditions, many populations show loss of these rhythms under constant darkness^36–41^. Cavefish populations have also convergently evolved a drastic reduction in sleep compared to surface fish^27^. Alterations can also be seen at the molecular level; an examination of clock genes (*per1, per2, cry1a*) in cave and surface fish found changes in the expression level and timing of activation of these genes in multiple cave populations.^36^ Consequently, the Mexican tetra provides a unique opportunity to study the evolutionary response of the biological clock and circadian rhythms to the arrhythmic cave environment across multiple cave populations.

In vertebrates, circadian rhythms are regulated by transcriptional feedback loops, where clock proteins directly or indirectly regulate the expression of the genes from which they are transcribed. The feedback loops of the circadian clock result in oscillations of gene expression of ~24 hours^42^. These oscillating transcripts make up the “circadian transcriptome” and are a substantial source of rhythmic physiology and behavior^43–45^. The conserved nature of the biological clock in vertebrates make comparative studies extremely powerful. Even across evolutionary timescales, many clock genes and downstream regulators of circadian behavior maintain similar functions.^45^ While many species of cave-dwelling organisms show evidence of weakened or absent locomotor rhythms^46–52^, no global analysis of transcriptional changes associated with clock evolution has been performed in any cave species, including *A. mexicanus*. Comparing global transcriptional shifts over circadian time between cave and surface populations can provide insight into naturally occurring dysregulation of the biological clock across multiple cave populations.

Here we characterize and compare the circadian transcriptome of *A. mexicanus* surface fish with that of three cave populations. We find widespread changes in the circadian transcriptome, with cave populations showing convergent reductions and losses of transcriptional oscillations and alterations in the phase of key circadian genes. Visualization of clock genes mRNAs of in the midbrain and liver of surface and cave fish support the dysregulation observed in the RNAseq data, and identify tissue-specific patterns unique to cave individuals. To functionally assess the contribution of dysregulated genes to sleep and behavior, we investigate the function of two genes linked to circadian regulation by creating CRISPR/cas9 mutants, and find that these genes are involved in regulating sleep behavior in *A. mexicanus*. Our results suggest that circadian rhythm, a highly conserved mechanism across most metazoans, is repeatedly disrupted at the molecular level in cavefish, with potential consequences for organismal physiology and behavior.

## Results

### Fewer genes show evidence of daily cycling in cavefish populations

To identify changes in rhythmic expression between cave and surface populations, we performed RNAseq with total RNA from whole fish from three cave populations (Molino, Pachón, and Tinaja) and one surface population (Río Choy) collected every 4-hours for one daily cycle under constant darkness at 30 days post-fertilization (dpf)(see Methods). Fry were raised in a light-dark cycle (14:10) and then transferred into total darkness 24 hours prior to the start of the sampling period. An average of 14,197,772 reads were mapped per sample (File S1, Fig. S1-4). Filtering of genes with low expression (< 100 total counts across all samples) resulted in 21,048 annotated genes used for downstream analysis. Although the cave populations are not monophyletic^53^, principle component analysis showed that the primary axis of differentiation among samples is habitat (*i.e.*, cave or surface; Fig. S5).

We used JTK_cycle^54^ to detect 24-hour oscillations in transcript abundance. We found that the surface population had the greatest number of rhythmic transcripts (539), followed by Tinaja (327), Pachón (88), and Molino (83), respectively (FDR < 0.05, see Table S1). Surprisingly few genes (19) showed significant cycling across all populations (Fig. 1A,B)(File S1). In all populations except Molino, we found significant overlap between rhythmic transcripts in *A. mexicanus* and those identified in zebrafish^55,56^ (See Methods; at *p* < 0.05: surface, Pachón and Tinaja, all *p* < 1 × 10^−4^; Molino *p* = 1, hypergeometric tests). Rhythmic transcripts in both the surface and cave populations were enriched for the GO terms related to circadian rhythm, including “regulation of circadian rhythm” (GO:0042752)(surface, *q* = 4.85 × 10^−10^; Tinaja, *q* = 4.16 × 10^−8^; Pachón, *q* = 2.33 × 10^−6^; Molino, *q* = 4.98 × 10^−6^). Rhythmic transcripts in the surface population were also strongly enriched for “visual perception” (GO:0007601)(*q* = 4.80 × 10^−12^), “sensory perception of light stimulus” (GO:0007602) (*q* = 9.28 × 10^−12^), and “phototransduction” (GO:0007602)(*q* = 1.14 × 10^−7^), unlike cave populations (lists of enriched terms in File S1).

**Figure 1.**
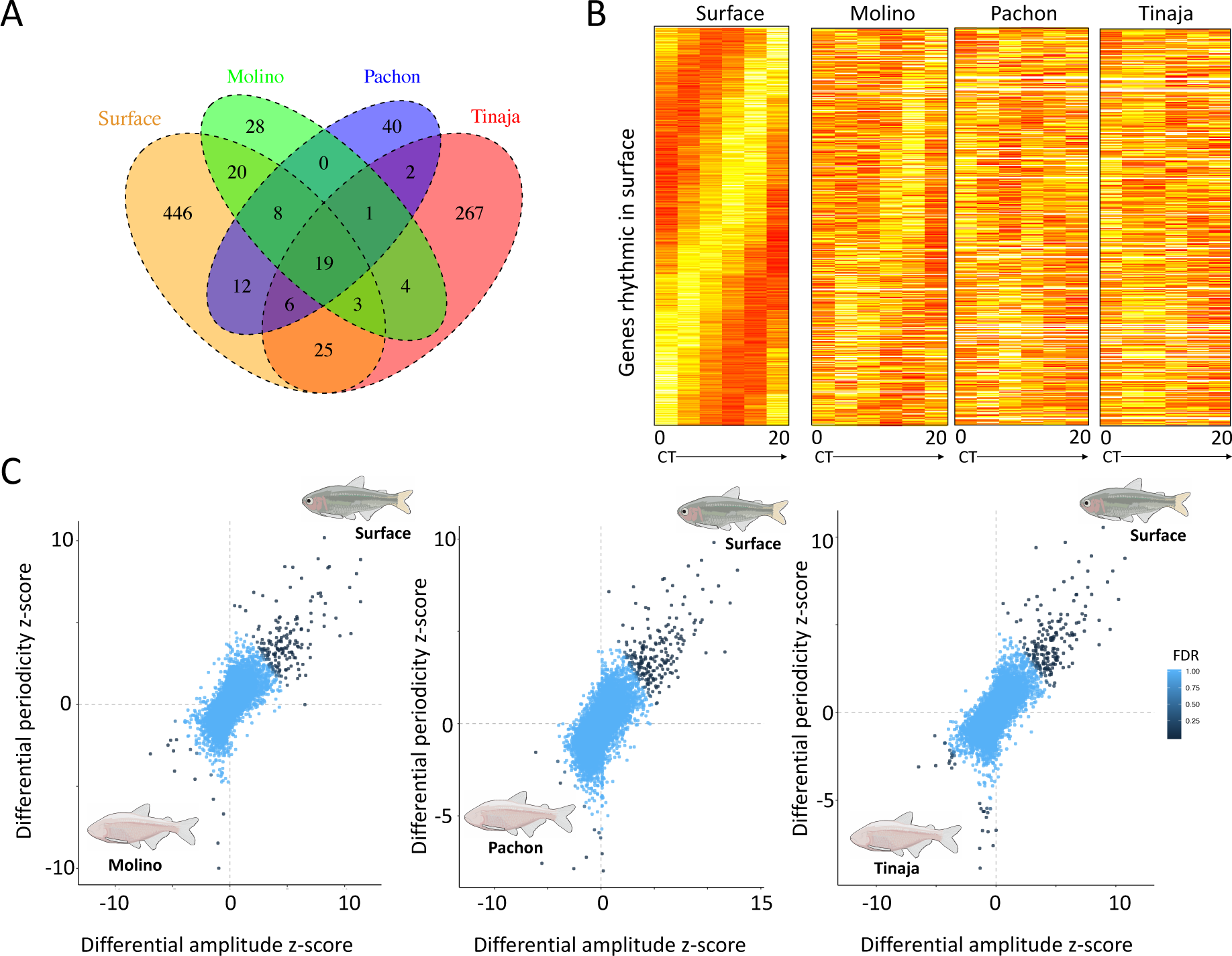
A. Overlap of genes with rhythmic expression between populations. B. Heatmap of genes with rhythmic patterns in surface fish (Río Choy), ordered by gene phase, compared to expression in cave populations. Each column represents gene expression at a single-time-point, sampled every four hours from 0-20 hours. Redder boxes correspond to higher expression. C. Identifying genes with changes in rhythmicity between cave and surface populations. Genes with greater amplitude values have larger differences between their expression peak and trough, where genes with greater periodicity show stronger cyclical oscillation patterns (see Methods). Genes are colored based on their *S*DR *q*-values. Genes with positive values for both show increased rhythmicity in the surface population, where genes with negative values show increased rhythmicity in cave populations.

Nearly 22% (117) of genes found to be rhythmic in the surface population were arrhythmic (*p-*value > 0.5) in all three cave populations and 77% (416) were arrhythmic in at least one cave population (Table S2, File S1)(see Methods), highlighting that the loss of rhythmicity has evolved independently among independent origins of the cave phenotype. Conversely, no genes that were rhythmic across all caves were arrhythmic in the surface population. Genes that were rhythmic in the surface population but arrhythmic in cavefish populations include genes known to be involved in the regulation of circadian rhythm (Table S3). For example, *arntl2* is a gene that encodes a transcriptional activator that forms a core component of the circadian clock and shows conserved rhythmic activity in zebrafish^55^ and mice^57^. Despite exhibiting a conserved function across vertebrates, this gene is arrhythmic in all three cave populations (surface, *p* = 0.0004, *q*=0.02; all caves, *p* > 0.5). Further, in comparing the *p-*values of genes with known roles in circadian rhythm (61 genes, see Methods, File S1), we found that these genes were significantly more rhythmic in their expression in the surface population than in cave populations (Wilcoxon signed rank test, *p* < 0.002 in all cave-surface pairwise comparisons).

### Cave and surface populations show differences in periodicity and amplitude of rhythmic transcripts

To identify genes with changes in rhythmicity between cave and surface populations, we calculated a differential rhythmicity score (*S*DR) for each gene for each population pair^58^. This metric accounts for both changes in how rhythmic a transcript is, as defined by differences in the JTK_cycle *p*-value for each gene between surface and cave populations, as well as differences in the robustness of a transcript’s oscillation, as defined by differences in amplitude of gene expression between the cave and surface populations. We found that 103 genes showed greater rhythmicity in the surface fish for all surface-cave comparisons (e.g., surface-Pachón, surface-Molino, surface-Tinaja; File S1)(Fig. 1C). This set of genes was highly enriched for the pathway “Circadian clock system,” with a 21-fold enrichment compared to the background set of genes used in this analysis (*q* = 9.96 × 10^−11^), and was the only pathway significantly enriched after false-discovery rate correction. Comparatively, relatively few genes showed significantly improved rhythmicity in cave populations (Fig. 1C, genes below zero for differential periodicity and differential amplitude z-score), and no genes showed increases in rhythmicity across multiple caves.

This unbiased assessment revealed that genes with reductions in rhythmicity in cave populations include several primary and accessory components of the core transcriptional clock (Fig. 2A)(Table 1). In the circadian clock’s primary feedback loop, members of the (bHLH)-PAS family (e.g., CLOCKs, ARNTLs) heterodimerize and bind to E-box DNA response elements to transcriptionally activate key clock proteins (e.g., PERs, CRYs). Negative feedback is then achieved by CRY:PER heterodimers which inhibit the CLOCK:ARNTL complex. Another regulatory loop is induced by the CLOCK:ARNTL complex activating the transcription of *nr1d1* and *rorc* genes (e.g., *rorca* and *rorcb*), which in turn both positively and negatively regulate the transcription of *arntl* (see Fig. 2B). Many of the genes that transcribe these activators and repressors show reductions or loss of cycling in one or more cave populations, and a few show reductions in rhythmicity in all three caves (e.g., *rorca, rorcb, arntl2*)(Table 1, Fig. 2). Reductions in rhythmicity at core clock genes suggest an overall dampening of the core circadian mechanism in cave populations compared to surface forms.

**Figure 2.**
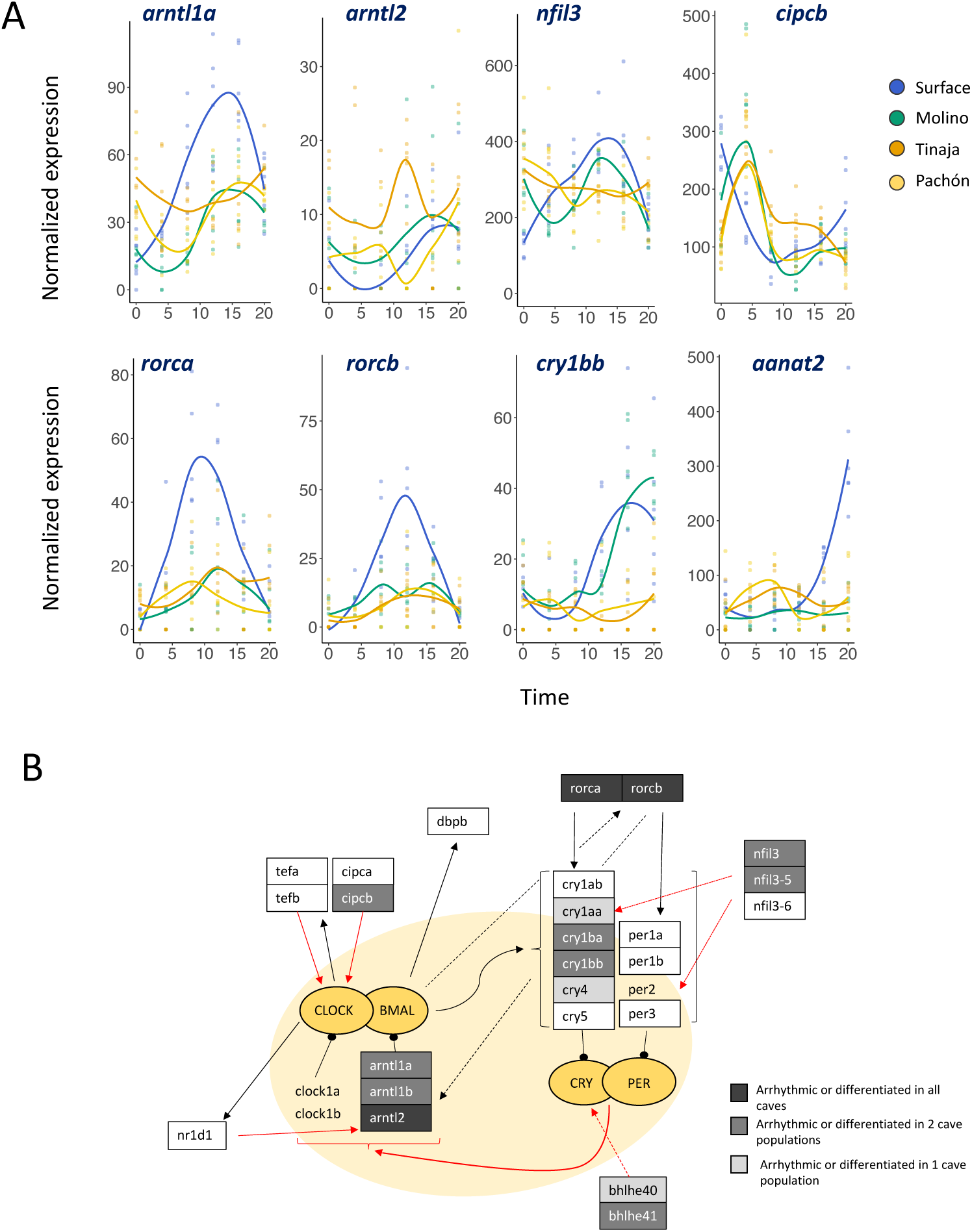
A. Key circadian genes with changes in rhythmicity in cave populations (see Fig. S17 for all core circadian genes with changes in rhythmicity). Colored lines represent a loess regression of gene expression through time for each population. B. Simplified schematic of the circadian feedback loops based on proposed interactions in zebrafish^43,45,55^. Grey boxes indicate genes that are either arrhythmic or show significantly reduced rhythmicity between cave and surface. White boxed genes do not show significant differences between cave and surface. Highlighted in yellow is the core loop. Bright yellow circles represent regulating protein complexes. Red lines indicate negative regulation, black lines indicate positive regulation. Genes that are not boxed did not show evidence of rhythmic expression in any cave or surface population. Dotted lines are for visual clarity. Genes without an annotated ortholog in cavefish were not included in the schematic.

**Table 1.**
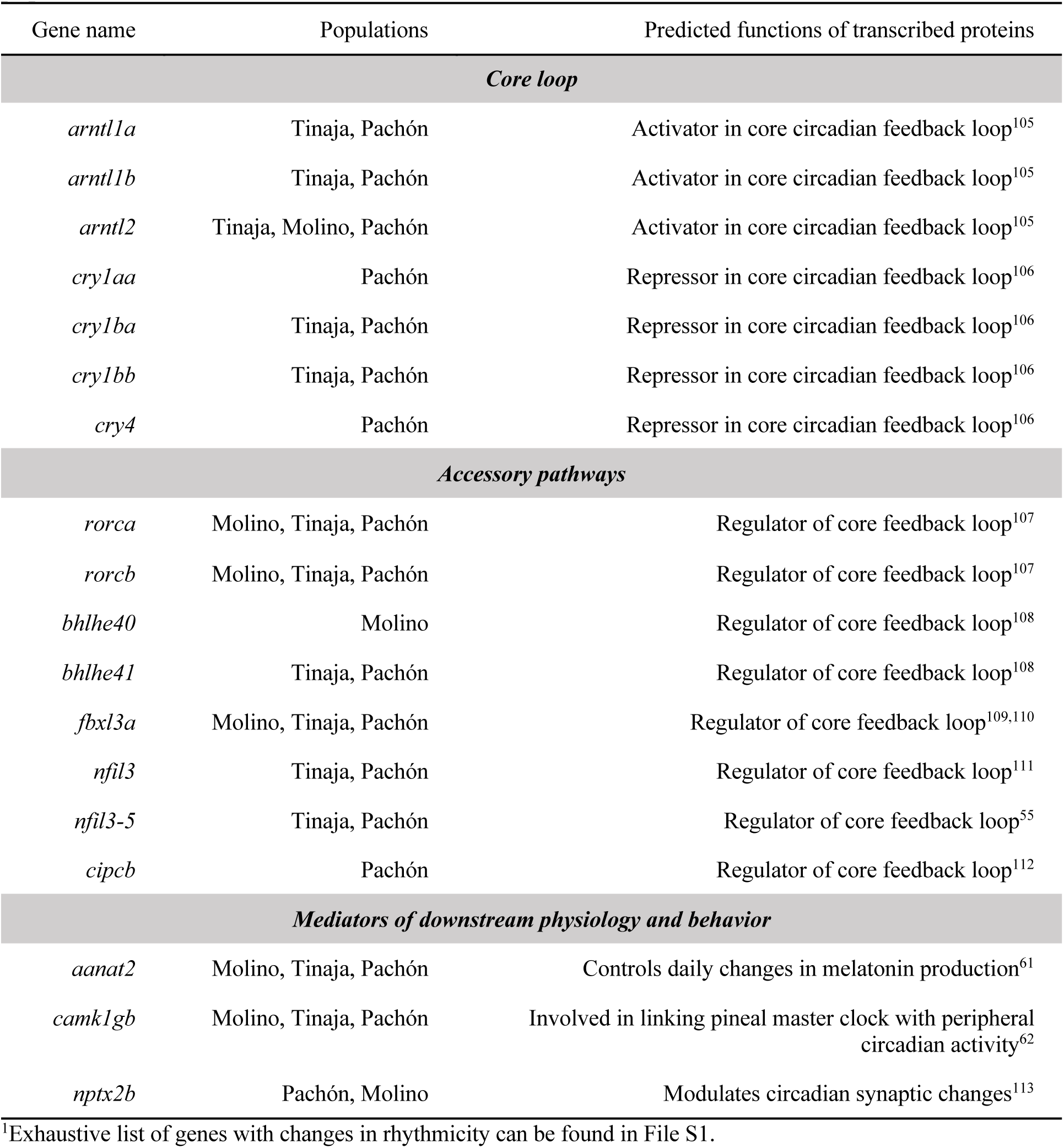
Known circadian regulators with losses or reductions in rhythmicity in cave populations^1^

| Gene name | Populations | Predicted functions of transcribed proteins |
| --- | --- | --- |
| *Core loop* |  |  |
| *arntl1a* | Tinaja, Pachón | Activator in core circadian feedback loop<sup>105</sup> |
| *arntl1b* | Tinaja, Pachón | Activator in core circadian feedback loop<sup>105</sup> |
| *arntl2* | Tinaja, Molino, Pachón | Activator in core circadian feedback loop<sup>105</sup> |
| *cry1aa* | Pachón | Repressor in core circadian feedback loop<sup>106</sup> |
| *cry1ba* | Tinaja, Pachón | Repressor in core circadian feedback loop<sup>106</sup> |
| *cry1bb* | Tinaja, Pachón | Repressor in core circadian feedback loop<sup>106</sup> |
| *cry4* | Pachón | Repressor in core circadian feedback loop<sup>106</sup> |
| *Accessory pathways* |  |  |
| *rorca* | Molino, Tinaja, Pachón | Regulator of core feedback loop<sup>107</sup> |
| *rorcb* | Molino, Tinaja, Pachón | Regulator of core feedback loop<sup>107</sup> |
| *bhlhe40* | Molino | Regulator of core feedback loop<sup>108</sup> |
| *bhlhe41* | Tinaja, Pachón | Regulator of core feedback loop<sup>108</sup> |
| *fbxl3a* | Molino, Tinaja, Pachón | Regulator of core feedback loop<sup>109,110</sup> |
| *nfil3* | Tinaja, Pachón | Regulator of core feedback loop<sup>111</sup> |
| *nfil3-5* | Tinaja, Pachón | Regulator of core feedback loop<sup>55</sup> |
| *cipcb* | Pachón | Regulator of core feedback loop<sup>112</sup> |
| *Mediators of downstream physiology and behavior* |  |  |
| *aanat2* | Molino, Tinaja, Pachón | Controls daily changes in melatonin production<sup>61</sup> |
| *camk1gb* | Molino, Tinaja, Pachón | Involved in linking pineal master clock with peripheral circadian activity<sup>62</sup> |
| *nptx2b* | Pachón, Molino | Modulates circadian synaptic changes<sup>113</sup> |

<sup>1</sup>Exhaustive list of genes with changes in rhythmicity can be found in File S1.

Genes with reduced rhythmicity in cavefish populations also include genes involved in downstream regulation of oscillations in physiology and behavior (Table 1). Among the genes which show reductions in rhythmicity across all three caves is *arylalkylamine N-acetyltransferase 2* (*aanat2*), which is involved in melatonin synthesis in the pineal gland^59–61^. *Aanat2* is required for the circadian regulation of sleep in zebrafish, and mutant zebrafish sleep dramatically less at night due to decreased bout length, but not decreased bout number^61^. Interestingly, cave populations show a dramatic reduction in sleep bout length compared to surface fish^27^. Like in zebrafish^59,61^, the expression of *aanat2* in surface *A. mexicanus* shows robust rhythmic behavior (*q* = 6.6 × 10^−4^) with peak expression during subjective night. In contrast, cavefish do not show evidence of rhythmic transcription of *aanat2* (Molino, *p =* 1.0; Tinaja, *p* = 0.82; Pachón, *p* = 0.33) and expression does not increase in these populations during subjective night (Fig. 2).

Another gene that shows reductions in rhythmicity across all three caves is *camk1gb*, a gene involved in linking the pineal master clock with downstream physiology of the pineal gland. Knock-downs of this gene in zebrafish significantly disrupt circadian rhythms of locomotor activity^62^, similar to what is observed in Pachón cavefish^39,40^. *Camk1gb* also plays a role in regulating *aanat2*; knockdowns of *camk1gb* reduce the amplitude of *aanat2* expression rhythm by half in zebrafish^62^.

### Rhythmic transcripts in cave populations show a delay in phase compared to the surface population

Next, we investigated whether the timing of the circadian clock differs between cave and surface populations by comparing the peak expression time (i.e., phase) of cycling transcripts. Most circadian genes show peak expression immediately before dawn or dusk^63^, and consistent with this, surface and cave populations of *A. mexicanus* all showed a bimodal distribution of phases (Fig. 3A). However, this distribution was shifted towards later in the day in fish from all three cave populations compared to the surface population. Our results are consistent with the individual gene analysis of Beale *et al.*^36^, which showed that the clock genes *per1* and *cry1a* display phase delays in fish from the Pachón and Chica caves relative to surface fish (Fig. S6). We then compared the phase of *A. mexicanus* genes with phase estimates of orthologous core clock genes in zebrafish sampled under dark conditions^55^. While zebrafish and *A. mexicanus* diverged ~146 MYA^64^, the timing of peak expression of core clock genes was highly similar in zebrafish and surface fish (median difference of 0.8 hours between orthologs, Table S4). Comparatively, the phase of clock genes in the core and accessory loops are often more shifted in cave populations relative to zebrafish phase than the surface population (median shifts of 4.7, 2.2, and 3.5 hours for Tinaja, Molino, and Pachón, respectively)(Table S4).

**Figure 3.**
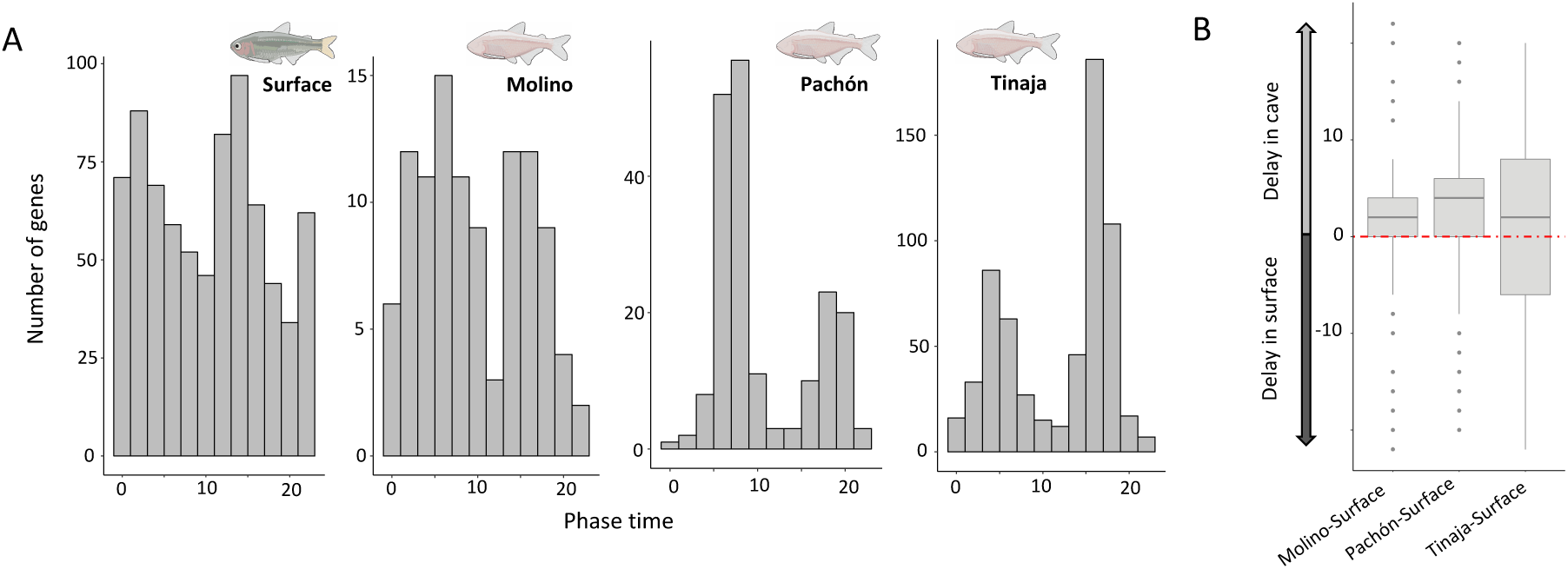
A. Phase (e.g., peak expression time) distribution of rhythmic transcripts in each population. B. Cycling genes on average show a delay in phase in cave populations relative to their phase in the surface population.

To further characterize differences in phase between surface and cave populations, we compared the phase of all genes with evidence for rhythmic expression across cave-surface population pairs (where *p* < 0.05 for a gene in the surface and the cave being compared). Consistent with the phase shifts observed in core clock genes and accessory loops, for each cave-surface comparison, we found that rhythmic transcripts showed significantly later peak expression in cave populations (Wilcoxon signed rank test, all pairwise comparisons *p <* 2.2 × 10^−16^, Table S5)(Fig. 3B, phase of all genes in File S1). Rhythmic transcripts in Pachón showed the greatest average shift, with a delay of 2.03 hours compared to surface fish (average shift, surface-Tinaja: 1.3 hours, surface-Molino: 0.48 hours).

### Genes with circadian cis- elements show loss of cycling and changes in phase in cavefish

In light of the alterations we see in the expression of genes in the circadian transcriptome, we next investigated circadian *cis-* elements in cavefish and surface fish. In vertebrates, circadian oscillations in gene expression are a consequence of interlocking transcriptional feedback loops of core clock proteins binding to a combination of known *cis-*elements (E-box, RRE, D-box) in the promoters and enhancers of target genes^55,65^. The timing of activator and repressor binding to these *cis-* elements results in the phase of oscillating transcripts^65^.

To better understand what drives oscillation patterns in *A. mexicanus*, we extracted promoter sequences of genes with rhythmic transcription in the surface population and identified transcription factor (TF) binding motifs in each sequence (Table S6). We consider rhythmic genes with significant circadian binding motifs (i.e., E-box, RRE, D-box sequences) putative targets of clock proteins in the circadian feedback loop. We then used a sliding window approach to identify the specific time window when the phases of circadian TF targets are most enriched in surface fish (see Methods). Consistent with observations in zebrafish^55^, the E-box, which is bound by CLOCK-ARNTL complexes as part of the core feedback loop^66^, showed phase-specific enrichment within the early peak (CT 2), and RRE elements, which are bound by RORA proteins and NR1D1/NR1D2 of the accessory loop^67^, were enriched within the later peak (CT 14-16)(Fig. S7)(see Methods). The D-box, which binds the repressor NFIL3, was also enriched within the early peak (CT 0-6), as in zebrafish^55^. The correspondence between surface *A. mexicanus* surface fish and zebrafish suggests that the timing of key regulatory cascades mediating circadian oscillations in gene expression is largely conserved between these species.

We then searched for circadian *cis-* elements in cavefish sequences upstream of genes that have lost rhythmicity in cave populations. A large proportion of the putative targets of core clock elements identified in the surface fish (e.g., genes identified as having an E-box, RRE or D-box motifs in their promoter sequence as described above) do not cycle in one or more cave populations: >77% are arrhythmic in at least one cave population. However, for nearly all gene for which we identified a proximal circadian motif in the surface fish, we also identified a binding site in cavefish (see Table S7 and Supplemental Methods). The maintenance of clock binding sites in cavefish populations suggests that the reduction in the number of rhythmic transcripts in cavefish is likely to be a consequence of changes in the transcription of core clock genes, rather than divergence at the target *cis-* binding sites at downstream genes.

We then compared the phase of putative clock targets in surface fish and cavefish. Consistent with the phase shifts seen in the core clock (Table S4), putative clock targets also show shifts in their phase in cavefish compared to surface fish (Table S8, Fig. S7). In the case of the E-box, in cave populations the phase intervals with the highest proportion of these motifs occur later (CT 4, 6, 6-10 in Molino, Pachón, and Tinaja, respectively) than what is seen in the surface population (CT 2) (Fig. S7). In contrast, for the RRE and D-box motif, there is no phase with enrichment in cave populations.

### Visualization of key circadian mRNAs reveals tissue-specific expression patterns in cave and surface populations

Our whole-fry RNAseq data suggest that rhythms in the core clock are dampened, and that rhythmic transcripts arephase shifted. Teleost circadian systems are highly decentralized and autonomous core clock gene expression may be observed in many tissues^68–70^, and tissue-specific differences may be obscured in whole organisms preparations^71^.

To address potential tissue-specific contributions to differences between cave and surface populations, we used RNAscope® fluorescence in situ hybridization (FISH)^72^. We collected brains and livers from individual 30dpf fry at 3 timepoints (CT0, CT8, and CT16)^70^ and performed RNA FISH for key circadian mRNAs in the primary loop (*per1* and *arntl1a*) and regulatory loop (*rorca* and *rorcb*) (Advanced Cell Diagnostics, Hayward, CA, USA, see Methods for details and controls). Brains were sectioned to allow visualization of the optic tectum and periglomerular grey zone (Fig. S8). These are both sites showing significant clock gene expression in zebrafish and these brain structures were consistently identifiable despite the small size of the 30dpf midbrain^73^.

We find that temporal expression patterns of *per1a* and *arntl1a* mRNA in the midbrains of surface fish is consistent with our RNAseq results and expression patterns in zebrafish^55^ (Fig. 4, ‘B’). *Per1a* expression peaks strongly at dawn (CT0) and rapidly diminishes. *Arntl1a* cycles in anti-phase and shows high expression at dusk (CT16). In contrast, cavefish show a number of population specific alterations to the temporal expression of *per1a* and *arntl1a* over the day (Fig. 4, S10, S11, S12). *Per1a* expression in cavefish shows broader temporal expression, consistent with phase shift at this gene seen in whole fish (Fig. S6). Where *per1a* expression rapidly diminishes in surface midbrain after CT0, expression of *per1a* persists through CT8 in cave populations. Surprising, Tinaja *per1a* expression is at its lowest at CT0, when the surface population *per1a* expression is greatest. Similarly, we see the persistence of *arntl1a* expression in Tinaja and Molino outside of CT16, the primary window of expression in the surface populations. Molino midbrains in particular showed high basal expression compared to the surface, specifically at CT8 when *per1a* and *arntl1a* expression is nearly absent in surface fish (Figs. S10, S12).

**Figure 4.**
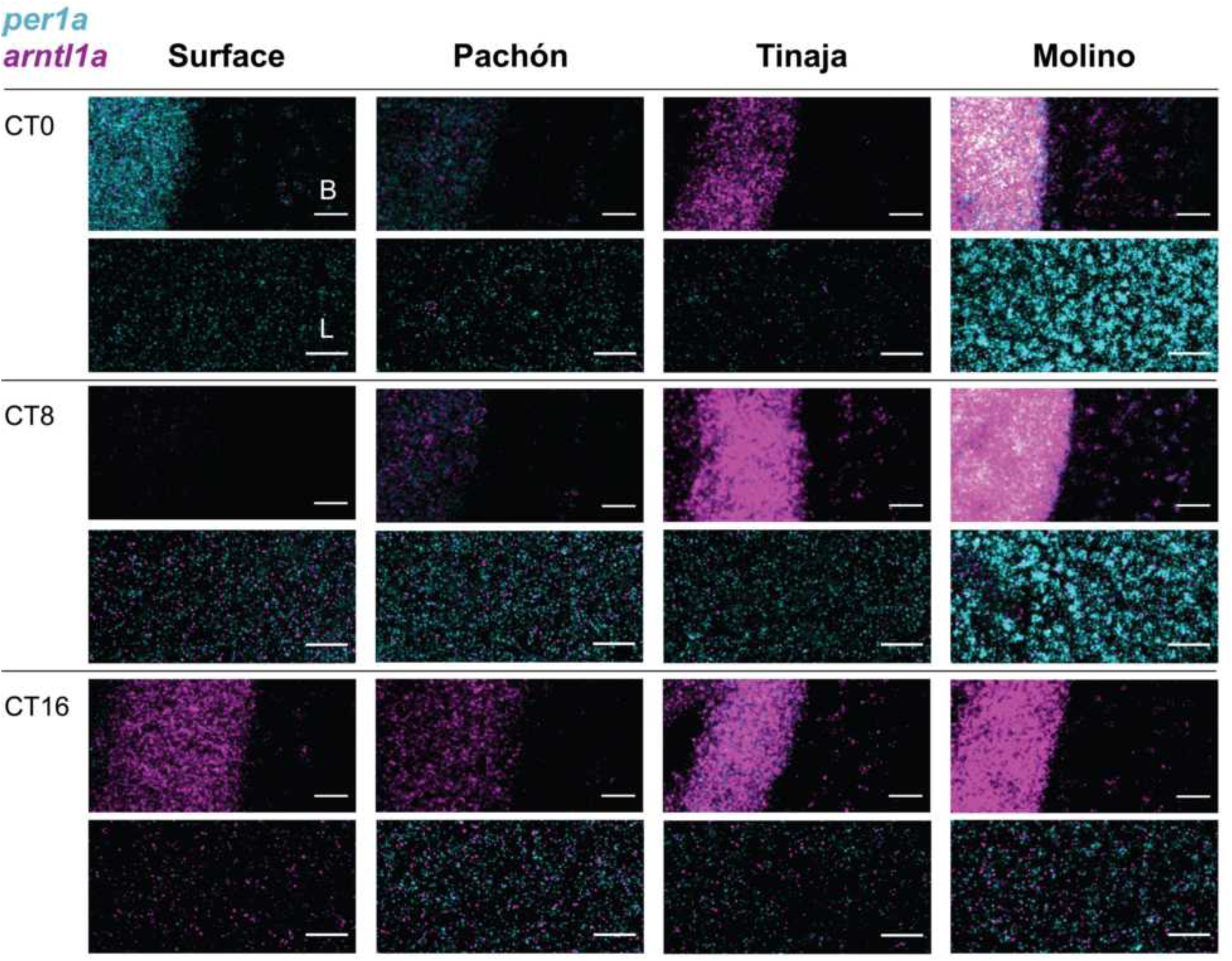
Temporal expression patterns of *per1a* and *arntl1a* in brain and liver tissue in *Astyanax mexicanus* populations. In-situ staining of *per1a* (cyan) and *arntl1a* (magenta) using RNAscope® in the midbrain (‘B’, top panels for each timepoint) and liver (‘L’, bottom panels for each timepoint) of surface fish and cavefish (Pachón, Tinaja, Molino) at CT0, CT8, and CT16. Each time point is a single fish sample with probes separated into two channels. Images are representative sections of two fish collected per time point, per population. Scale bar is 25μM.

Temporal expression patterns for *per1a* and *arntl1a* in surface fish is also similar to zebrafish (Fig. 4, ‘L’). Surface fish liver expression of *per1a* and *arntl1a* are largely in synchrony with the expression of these genes in surface midbrain, however, expression of *per1a* and *arntl1a* in the liver is also seen at the intermediate timepoint, CT8. Compared to surface, Pachón and Tinaja livers show similar *arntl1a* expression but peak expression of *per1a* is delayed (occurring at CT8 instead of CT0). *Per1a* expression persists in Pachón and Tinaja at CT16, when *per1a* transcripts are low in surface livers. Intriguingly, peak and trough expression in Pachón and Tinaja livers appear to be out of sync with those in the midbrains (Fig. 4, ‘L’ vs. ‘B’). *Per1a* expression in Tinaja liver is greatest at CT8, but is expressed highest in brains at CT16. In Pachón midbrains, *per1a* expression is highest at CT0, but expression is highest in the livers at CT8 and CT16. In contrast, Molino livers exhibit temporal expression of *per1a* that is synchronous with brains. However, as with the midbrain, *per1a* has high basal expression in Molino compared to surface, Pachón, and Tinaja.

Next, we examined the expression pattern and timing of *rorca* and *rorcb*, two gene members of the regulatory loop that regulate the expression of *arntl* paralogs and circadian transcription^67^. Our RNAseq results indicate that *rorca* and *rorcb* show peak expression in whole surface fish at CT12 (Fig. 5, S13), similar to what has been observed in zebrafish (Table S4). Consistent with this, in RNA FISH of surface fish livers and midbrains, *rorca* and *rorcb* expression was highest midday (Fig. 5, S13). Temporal expression of *rorca* and *rorcb* was similar between livers and the midbrain in cave populations. When compared to surface tissues, the expression of *rorca* and *rorcb* is delayed in Pachón and much broader in Tinaja and Molino populations. Molino midbrains have high expression of *rorca* and *rorcb* compared to all other populations. In addition, *rorcb* expression is seen across timepoints in all caves, particularly in Molino, whereas it is nearly absent in surface fish at CT0 and CT8 (Fig. 5).

**Figure 5.**
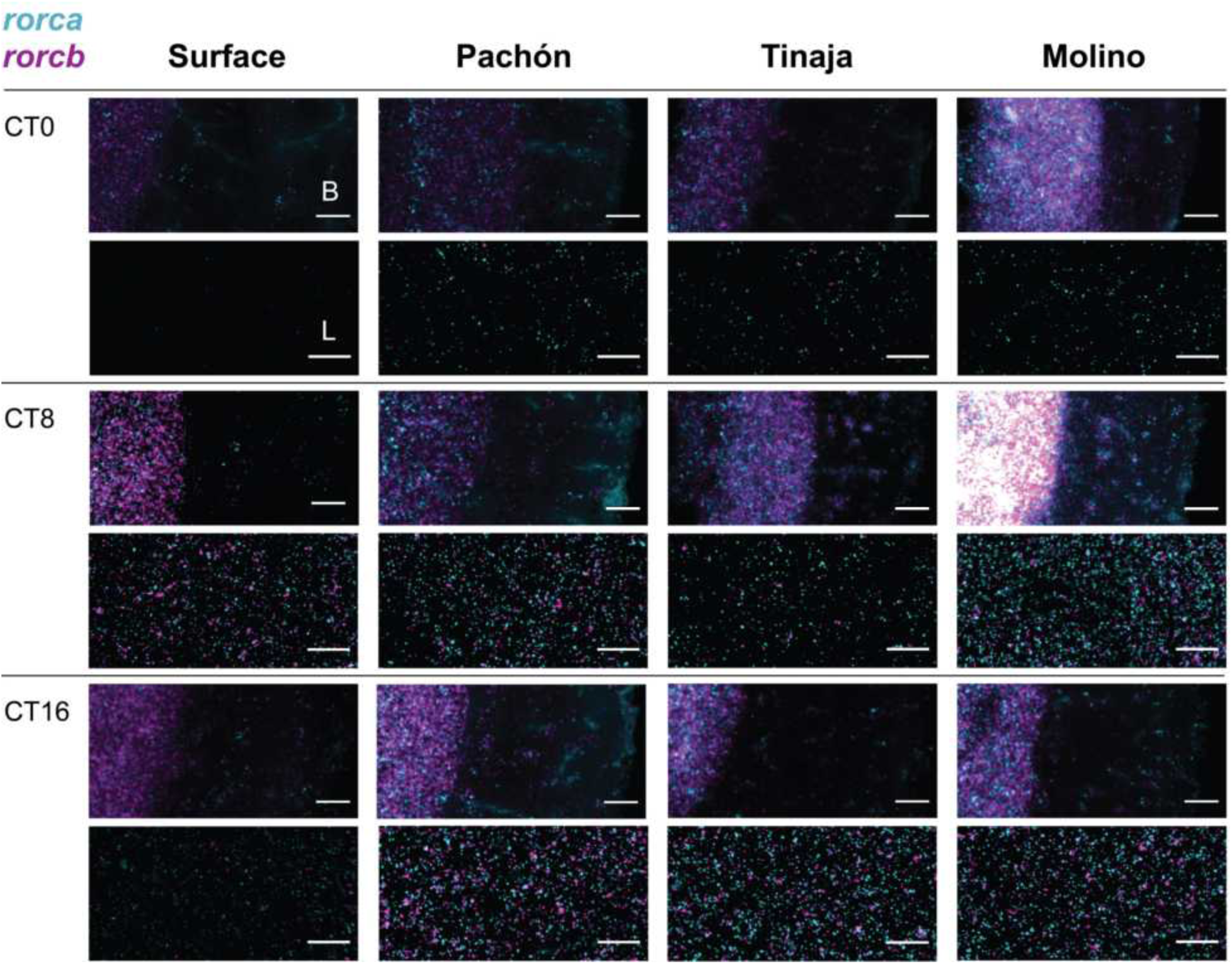
Temporal expression patterns of *rorca* and *rorcb* in midbrain and liver tissue in *Astyanax mexicanus* populations. In-situ staining of *rorca* (cyan) and *rorcb* (magenta) using RNAscope® in midbrain (‘B’, top panels for each timepoint) and liver (‘L’, bottom panels for each timepoint) of Surface fish and cavefish (Pachón, Tinaja, Molino) at CT0, CT8, and CT16. Each time point is a single fish sample. Images are representative sections of two fish collected per time point, per population. Scale bar is 25μM.

Thus, as observed in the whole organism RNAseq analysis, tissue-specific expression of these primary loop and regulatory loop genes confirm the phase shifts and dysregulation of cavefish transcripts compared to surface fish, though, our RNAscope analysis suggests that disruptions to the temporal regulation *per1a* and *arntl1a* expression may not manifest uniformly across tissues in Pachón and Tinaja cavefish (Fig. 4).

### *Aanat2* and *rorca* regulate sleep behavior in surface fish

The circadian clock plays an important role in regulating the sleep-wake cycle^70^. We have found that expression of the genes in the core pacemaker, as well as a circadian sleep regulator (*aanat2*), are differentiated between cave and surface populations (see Fig. 2). Cavefish populations have repeatedly evolved towards a reduction in sleep at night and overall, raising the question of whether dysregulation of the biological clock is contributing to changes in sleep behavior between cave and surface populations. However, the function of circadian genes in *A. mexicanus* sleep regulation has not been investigated.

To explore the role of clock genes in sleep regulation in *A. mexicanus*, we created crispant surface fish mosaic for mutations in *aanat2* and *rorca*. *Rorca* and *aanat2* were selected because they show differences in expression between the surface and all cave populations, and because *aanat2* plays an important role sleep regulation in zebrafish^61^. Mutagenized 30 dpf fish and wildtype (WT) sibling controls were phenotyped for sleep behavior. Raw locomotor behavior was processed to quantify sleep duration and behavioral architecture (locomotor distance, waking activity, sleep bout duration, and sleep bout number) as previously described^38,74^. For each injected construct, three separate biological replicates were carried out to ensure the phenotype was consistently reproducible. Genotyping confirmed mutagenesis (Figs. S14, S15).

*Aanat2* crispants demonstrate the role of this gene in the regulation of sleep in *A. mexicanus*. While there was no difference in total sleep duration between WT surface and *aanat2* surface fish crispants (unpaired t-test, *p* = 0.98, Fig. 6A), night sleep duration (minutes/hour) was significantly reduced in *aanat2* crispants (unpaired t-test, *p* = 0.0108, Fig. 6B,C). These results are consistent with the function of this gene in zebrafish, where increased expression of this gene during subjective night increases melatonin production, regulating nighttime sleep^61^. Zebrafish *aanat2* knockout mutants produce little or no melatonin and sleep almost half as much during the night as control fish^61^.

**Figure 6.**
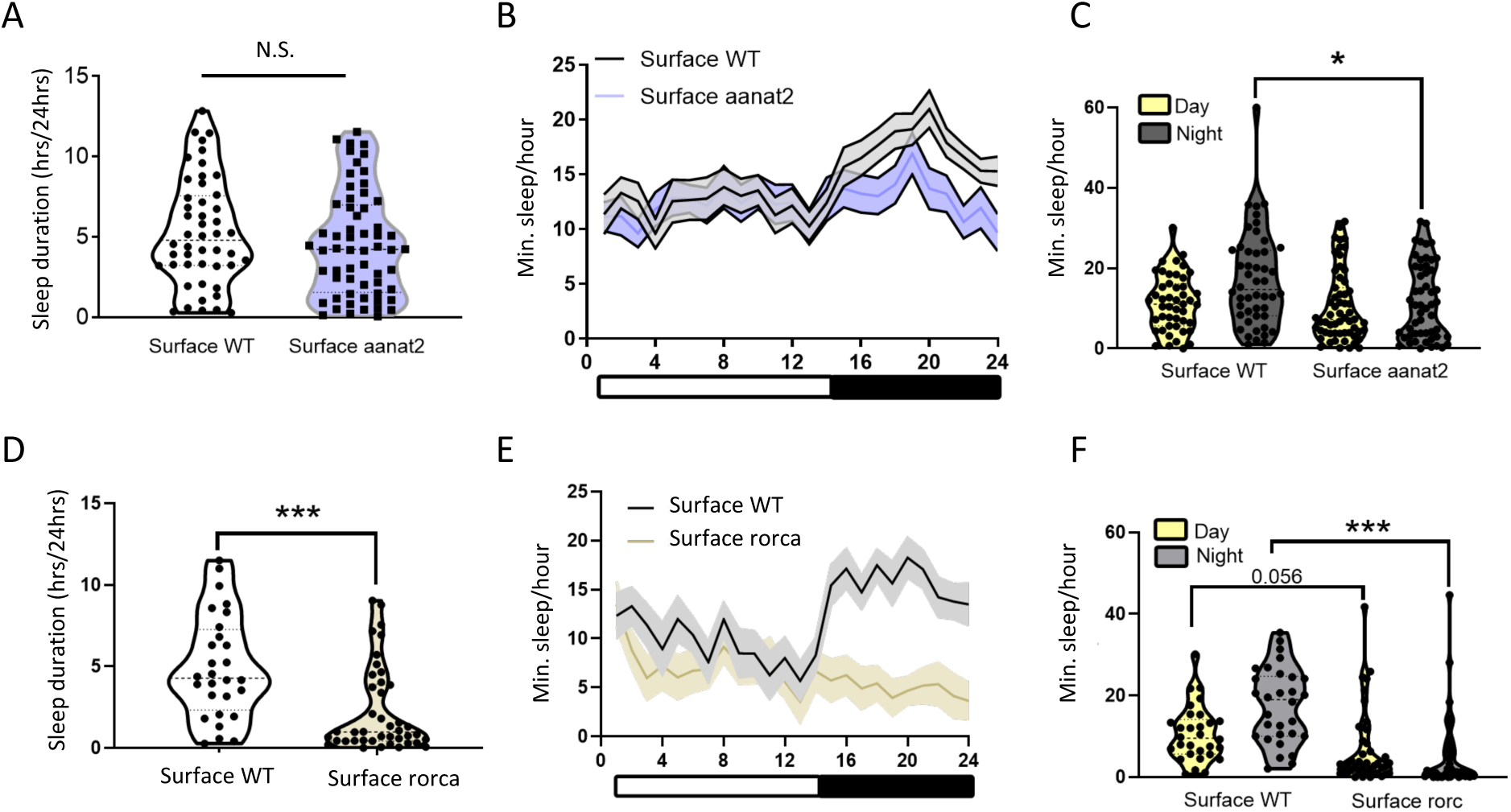
Mutant *aanat2* and *rorca* fish reveal a role for these genes in sleep behavior in *A. mexicanus*. **A**. Total sleep is not significantly altered between control and *aanat2* crispant surface fish (unpaired t-test, *p*=0.99). **B-C.** Day sleep is not significantly altered between WT and crispant *aanat2* fish (unpaired t-test, *p*=0.55). Night sleep is significantly reduced in *aanat2* crispants compared to WT controls (unpaired t-test, *p*<0.0001). **C.** Total sleep is significantly reduced in crispant *rorca* fish compared to WT controls (unpaired t-test, *p*<0.0001). **E-F.** Day sleep was not significantly altered between WT and *rorca* crispants (unpaired t-test, *p*=0.056). Night sleep is significantly reduced in *rorca* crispants compared to WT controls (unpaired t-test, *p*<0.0001).

*Rorca* crispants also show differences in sleep behavior compared to WT surface fish. Like *aanat2* crispants, *rorca* mutants sleep significantly less minutes per hour at night compared to WT sibling fish (unpaired t-test, *p* < 0.0001, Fig. 6E,F); however, *rorca* crispants also sleep less overall during a 24-hr period (unpaired t-test, *p* < 0.0001, Fig. 6D). While WT surface fish sleep an average of 5.16 hours per 24-hr period, *rorca* surface mutants sleep just 2.26 hours per 24-hr period. *Rorca* expression has not been previously associated with sleep regulation, but it is a transcriptional regulator of the core clock gene *arntl1a/b*. The mammalian ortholog of *arntl1a/b, arntl1*, has been implicated as a regulator in the sleep-wake cycle and deletions alter sleep behavior in mammals^75–77^. Altogether, these results demonstrate that alterations to the biological clock in surface fish can alter sleep behavior. Future investigations into the specific genetic variants between surface and cavefish will help delimit the role of the biological clock in cavefish sleep phenotypes.

## Discussion

Circadian rhythms are nearly ubiquitous in eukaryotes, directing aspects of behavior, physiology, and metabolism^2^. The biological clock is thought to provide a mechanism for organisms to synchronize their physiology and behavior to predictable daily cycles^3,4^. However, we have a limited understanding of how the biological clock is altered in the face of arrhythmic environments, where organisms are isolated from regular environmental cues^15,36,42,78,79^. Caves provide a natural laboratory for perturbations to circadian biology: isolated from light-dark cycles and with little fluctuation in temperature, cave-dwelling organisms lack many of the environmental stimuli found outside caves^15,16,36^.

Here we have utilized comparisons between conspecific *Astyanax mexicanus* cavefish and surface fish to explore changes to the biological clock following adaptation to the cave environment. Using RNAseq to survey gene expression changes over a 24-hr period, we identified several repeated alterations in the biological clock in cavefish populations, including: 1) a reduction in the number of transcripts with evidence of daily cycling, 2) the changes in the oscillation patterns of genes in the primary circadian feedback loop (e.g., *arntl2*, *arntl1a/b*, *cry1ba, cry1bb*) and accessory loops (*rorca/b*, *bhlhe40/1*), and 3) an overall delay in phase of the genes in the circadian transcriptome, including core clock genes and their putative targets. Population genetic analyses indicate that a number of genes in the circadian transcriptome are also genetically differentiated between the surface and one or more cave populations (see Supplemental Material and Methods). Visualization of mRNAs of core (*arntl1a*, *per1a*) and accessory loop (*rorca*, *rorcb*) in the midbrain and the liver of cave and surface fish support changes in the regulation of key clock genes and highlight tissue-specific changes in the expression of core clock genes in two of the cavefish populations relative to surface fish. The decoupling of rhythms between tissues in caves is an interesting biological feature that warrants further investigation across more tissues in these populations.

In addition to changes in the core and accessory loops, we also found that an important regulator of circadian physiology, *aanat2*, shows reductions in rhythmicity across all three cave populations. *Aanat2* encodes a protein that regulates melatonin production in the pineal gland, and robust expression of this gene is considered a marker for circadian clock function^59,61,80,81^. We found that cavefish populations, unlike surface fish and other teleosts^82^, did not show increased *aanat2* expression during the subjective night (Fig. 2), making dysregulation of the circadian clock an interesting candidate for the repeated changes in sleep regulation in seen across cavefish populations^27^. Investigating the function of *aanat2* and *rorca* through the creation of *aanat2* and *rorca* crispants revealed that these genes play a role in sleep regulation in *A. mexicanus*. Consequently, we speculate that dysregulation of the biological clock may have behavioral consequences for cavefish, and in particular, may contribute to the reductions in sleep seen across cavefish populations.

Beyond the regulation of circadian physiology, environmental light plays an essential role in cell cycle control and activating DNA repair in teleosts^83–86^. Previous work in this system has found that cave populations of *A. mexicanus* show higher levels of DNA repair activity and increased expression of DNA repair genes in the dark^36^. As a common light signaling pathway controls both DNA repair and clock entrainment, one purposed mechanism for this is that cavefish have sustained upregulation of clock genes normally driven by light, akin to ‘perceiving’ sustained light exposure^36^. While we found that DNA repair genes are more often upregulated in cave populations (Table S10, see Supplemental Material and Methods), we did not find that genes associated with light induction were more likely to be upregulated in cavefish compared to surface fish under dark-dark conditions (Table S11). Nor did we find that light-activated genes in the core circadian feedback loop are consistently upregulated in cave populations (see Supplemental Material and Methods, Figure S16). Thus, more work is necessary to understand the relationship between clock evolution and DNA repair in this system.

Altogether, these results suggest that highly conserved features of circadian biology have been repeatedly disrupted across populations of cavefish. Further research on the molecular basis of clock dysregulation will help delineate if clock dysfunction is a consequence of molecular alterations to (1) the pathways that perceive, or synchronize the biological clock to, external stimuli, or (2) alterations to the core transcriptional feedback loop, and whether this dysregulation is a consequence of selection or drift after the movement into the cave environment.

## Methods

### Sampling

Samples from each population were derived from the same mating clutch and raised in 14:10 light-dark cycle. All fish were exactly 30 days post fertilization at the start of the experiment. Fish were kept in total darkness for 24 hours prior to sampling and throughout the duration of the experiment. Six replicates were first sampled at 6am (Circadian Time 0) and then every four hours throughout until 2am (corresponding to CT 20). During the experiment, fish were fed *ad libitum* twice daily, in the morning and evening (exactly at 8am and 8pm). Individuals were immediately flash frozen in liquid nitrogen. RNA was extracted from the whole organism with extraction batch randomized for time of sampling and population. Samples were sequenced on the Illumina HiSeq 2500 platform to produce 125-bp paired-end reads (File S1) using strand-specific library preparations. Samples were randomized across lanes relative to population and time of sampling. Previous work showed no extraction batch or lane effect on these samples^87^.

### Read mapping

Reads were cleaned of adapter contamination and low-quality bases with Trimmomatic (v0.33)^88^. Cleaned reads were mapped to the *A. mexicanus* draft genome assembly v1.02 (GCA_000372685.1) with STAR^89^ and reads overlapping exonic regions were counted with Stringtie (v1.3.3d)^90^ based on the *A. mexicanus* Ensembl v91 annotation. Genes with fewer than 100 reads across all samples were removed from the analysis. Reads were subsequently normalized for library size and transformed with a variance stabilizing transformation with DESeq2^91^ for principle component analysis.

### Identification of rhythmic transcripts

Rhythmic transcripts with 24-h periodicity were identified using the program JTK_cycle^54^, using one 24-h cycle, with a spacing of 4 hours and 6 replicates per time point. JTK_cycle was also used to estimate the amplitude and phase of each rhythmic transcript. We retained genes as rhythmic at an False Discovery Rate (FDR) of 5%. Our overall result, that more genes are cycling in the surface population that in cave populations (Table S1), was insensitive to specific value of the FDR cut-off. We defined arrhythmic transcripts at *p*-value > 0.5, a conservative cut-off for arrhythmic expression that has been used in other studies^58,92^.

### Differential rhythmicity analysis

To identify differential rhythmicity between pairs of populations, we calculated a differential rhythmicity score (SDR) for genes with a JTK_cycle *p*-value < 1 in either population. As in Kuintzle *et al.*^58^, SDR was defined as:

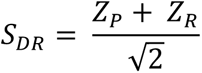

Where Zp is the Z-score for changes in periodicity between populations, where changes in periodicity are defined as log(*p* -value of population 1) – log(*p* -value of population 2). ZR is the Z-score computed for changes in amplitude between populations, where changes in amplitude are defined as: log2(Amplitude of population 2 / Amplitude of population 1). Amplitude for each gene was estimated with JTK_cycle.

We then computed a *p*-value for SDR values using a Gaussian distribution based on the fit to the empirical distribution. We used R’s *p.adjust* to perform a Benjamini & Hochberg correction for multiple testing (Table S9).

### Promoter analysis of rhythmic genes

We defined putative promoters as the region 1-kb upstream to 200bp downstream from the transcription start site (TSS). Population genetic data^53^ was used to identify variants with putative promoter regions segregating between populations and create alternative reference genomes for each population. Genes with a JTK_cycle FDR < 0.1 in the surface were used to identify phase enrichment of circadian *cis-* elements in the surface population. FIMO of the MEME suite^93^ was used to identify motifs in each promoter. FIMO was used to estimate *p*-values for motifs in each sequence; motifs with *p*-values < 0.0001 (the default cut-off) were considered for downstream analyses. To identify phase-specific enrichment of binding motifs, we tested sliding windows of time with Fisher’s exact tests for motifs in phase versus motifs out of phase compared to the total set of in- and out-of phase genes. To compute changes in phase of genes that are putative targets of circadian transcription factors between surface and cave populations, we calculated the minimum distance between the two phases based on a 24-h clock. Further details of this analysis are available in the Supplemental Methods.

### Tissue preparation for RNA fluorescence in situ hybridization (FISH)

Pachón, Molino, Tinaja, and surface fish derived from the Río Choy population were raised in densities of 20-25 individuals/3L tank in 14:10LD cycle at 23°C and 750-800μS/cm conductivity, with feeding and water quality control as previously described^94^.

29dpf fish were placed in constant darkness in the dark phase of the day preceding tissue collection. 30dpf fish were fed *ad libitum* at 8am and 8pm on the day of tissue collection. Individual fish were euthanized in MS-222 at CT0, CT8, and CT16. Experiments were performed in duplicate: two fish were euthanized and dissected at each time point from each population. All efforts were made to reduce light exposure prior to fixation. Dissected brains and livers were placed into a freshly prepared solution of 4% PFA in DEPC 1X PBS on ice. Samples were transferred to room temperature after 3-5 minutes on ice and allowed to fix for 1 hour. Samples were then placed at 4°C for 36 hours. After 36 hours, samples were rinsed briefly in DEPC water and washed in DEPC 1X PBS three times for 15 minutes with constant shaking. A graded EtOH wash was performed (30%- 50%- 70%- 100%) in DEPC treated water with 5 minutes between steps.

Tissue processing and paraffin embedding were performed with a PATHOS Delta hybrid tissue processor (Milestone Medical Technologies, Inc, MI, USA)^95^. Paraffin sections were cut with 12μm thickness using a Leica RM2255 microtome (Leica Biosystems Inc. Buffalo Grove, IL, USA) under RNase free conditions. Brains were sectioned coronally through the mesencephalon-diencephalon to allow visualization of the optic tectum and periventricular grey zone, sites showing significant clock gene expression in zebrafish^73^ (Fig. S8). We were unable to consistently section through the same region of the brain in order to compare expression in hypothalamic nuclei due to the size of 30dpf brains^73^. Livers were sectioned longitudinally. Sections were mounted on SureBond charged microscope slides (cat#SL6332-1, Avantik, Springfield, NJ, USA). To allow for comparison between populations, all brains or livers from a single probe set were processed at once (e.g., *rorca-rorcb* brains from all populations, and all time points were processed at the same time). In addition, brains or livers from each time point (CT0, CT8, CT16) were processed on a single slide to allow for a direct comparison between time points within populations.

### RNA FISH, imaging and analysis

RNAscope® probes were designed by ACDBio to target all known mRNA transcripts of genes *per1a* (ENSAMXG00000019909), *arntl1a* (ENSAMXG00000011758), *rorca* (ENSAMXG00000009363), and *rorcb* (ENSAMXG00000015029) (Advanced Cell Diagnostics, Hayward, CA, USA) based on Astyanax_mexicanus-2.0, INSDC Assembly (GCA_000372685). Probes were ordered for two-plexing (*per1a-arntl1a* and *rorca-rorcb*) to provide an internal control in each sample for mRNA phase and quantity. Probes may be ordered at the ACD Online Store using the following catalog numbers: *per1a* (590801), *arntl1a* (590831-C2), *rorca* (590811), and *rorcb* (590821-C2). *In situ* hybridization of *per1a, arntl1a*, *rorca*, and *rorcb* mRNA was performed on paraffin sections using RNAscope Multiplex Fluorescent Detection Kit v2 (Advanced Cell Diagnostics, Hayward, CA, USA) according to the manufacturer’s protocol.

Slides were stored in the dark at 4°C before imaging. Fluorescent images of sections were taken with a Nikon Eclipse TI equipped with a Yokogawa CSU W1 spinning disk head and Hamamatsu ORCA-Flash 4.O camera for high-resolution imaging using a Nikon 20x/0.75 Plan Apo objective. Probes *rorca* and *per1a* were imaged with 561nm laser and collected with an ET605/70m emission filter, and *arntl1a* and *rorcb* with 633nm laser and collected with an ET700/75m emission filter. DAPI was excited at 405nm and collected with an ET455/50m emission filter. Samples were identified automatically for imaging using a custom script as in Guo *et al*.^96^, and each sample was imaged as a tile scan with 10% overlap and z-stack of depth of 18µm with 0.9 µm optical slice thickness. Microscope and camera settings during acquisition were identical across all samples to allow for a direct comparison between timepoints and populations. Image processing in Fiji^97^ was identical between populations and time points within a tissue and probe set. Briefly, tiles were stitched into a complete image using Grid/Collection Stitching^98^, maximum projected and contrast adjusted. To properly compare cave populations to surface populations, we set the intensity ranges for each image to that of the surface sample images (Fig. 4). The intensities of Molino and Tinaja brain images in Fig. S11 are adjusted for oversaturation. Figs. 4 and 5 exclude the DAPI channel. Corresponding DAPI staining for these figures can be found in Fig. S9. Images are representative of two fish collected from each timepoint per population. For Figs. 4 and 5, maximum projected images are shown.

For quantification in Fig. S10, tiles were stitched into a complete image using Grid/Collection Stitching^98^. Stitched images were sum projected and background subtracted. 400×400 anatomical regions of interest were identified by visual inspection and average intensities were measured for each channel. To allow for a comparison of relative expression per cell from multiple samples, DAPI intensity was measured as a proxy for cell density, and FISH signal intensity was normalized against DAPI intensity for each region. Technical replicates (two to three sections from the same liver or brain) were averaged to make biological replicates represented as points in Fig. S10. All groups have two biological replicates with the exception of *rorca*-*rorcb* Pachón brains which have only one due to sample loss. Graphing and statistical analysis were performed using GraphPad Prism software (GraphPad, Prism version 8.3.0, GraphPad Software, San Diego, California USA). For each mRNA probe, cave population means were compared with the control mean (Surface) within timepoints using 2-way ANOVA. Dunnett’s test was used to correct for multiple comparisons across populations and timepoints. Original data underlying this part of the manuscript can be accessed from the Stowers Original Data Repository at http://www.stowers.org/research/publications/libpb-1485.

### CRISPR/Cas9 design and genotyping

Functional experiments were performed by generating crispant fish mosaic for mutations in *aanat2* and *rorca*. CRISPR gRNAs were designed using ChopChop v3 software^99^. gRNAs were designed to target an exon in each of the genes and to produce a double stranded break close to a restriction enzyme site for genotyping purposes. For *aanat2*, the gRNA was 5’-GGTGTGCCGCC<u>GCTGCC</u>GGA-3’ and for *rorca* the gRNA was 5’-gGAGAACGGTAACG<u>GCGGGC</u>A-3’ where the lower case g at the 5’ end was added to the sequence for T7 transcription (restriction enzyme target sites are underlined). gRNAs were synthesized as previously described^100^ with modifications^101,102^. Briefly, gRNA specific oligos were synthesized (IDT) that contained the gRNA target site, T7 promoter, and an overlap sequence:

> *aanat2* oligo A:
>
> 5’- TAATACGACTCACTATAGGTGTGCCGCCGCTGCCGGAGTTTTAGAGCTAGAAATAGC-3’
>
> *rorca* oligo A:
>
> 5’- TAATACGACTCACTATAgGAGAACGGTAACGGCGGGCAGTTTTAGAGCTAGAAATAGC-3’
>
> Each oligo A was annealed to the oligo B:
>
> 5’- GATCCGCACCGACTCGGTGCCACTTTTTCAAGTTGATAACGGACTAGCCTTATTTTAACTTGCTATT TCTAGCTCTAAAAC-3’

and these annealed oligos were amplified. gRNAs were transcribed using the T7 Megascript kit (Ambion), as in Klaassen *et al.*^103^ and Stahl *et al.*^102^, and purified using a miRNeasy mini kit (Qiagen). Nls-Cas9-nls^104^ mRNA was transcribed using the mMessage mMachine T3 kit (Life Technologies) following the manufacturer’s instructions and purified using the RNeasy MinElute kit (Qiagen) following manufacturer’s instructions. 150 pg Cas9 mRNA and 25 pg *aanat2* gRNA or 150 pg Cas9 mRNA and 25 pg *rorca* gRNA were injected into single-cell surface fish embryos (2 nL/embryo were injected).

Genomic DNA from injected embryos and wild-type (uninjected) controls was extracted at 48 hours post-fertilization^101,102^ and used for genotyping by PCR to determine if mutagenesis was achieved at the locus through gel electrophoresis (Figs. S14, S15). Briefly, gRNAs were designed to disrupt specific restriction enzyme sites; samples were incubated with that restriction enzyme to determine if the Cas9 cut. The undigested fragment acts as a control while the digested fragment is used to genotype. Cases where we see a band where the undigested fragment is indicates that the restriction enzyme was not able to cut due to the site being disrupted because of the CRISPR/Cas9 + gRNA. Gene specific primers are given in File S1. PCRs were performed with a 56 °C annealing temperature and a 1-minute extension time for 35 cycles. The resulting PCR product was split in half, and one half was restriction enzyme digested. A DNA fragment containing the *aanat2* target fragment was digested with BbvI (New England Biolabs Inc.). The *rorca* DNA fragment was digested with Cac8I (New England Biolabs). Both DNA fragments were digested at 37 °C for 1 hour.

For wildtype individuals, the PCR fragment was used for cloning. For crispant fish, restriction enzyme digests were performed (as described above) and the uncleaved bands were gel purified using a gel extraction kit (Qiagen). PCR products were cloned into the pGEM®-T Easy Vector (Promega), and three clones per individual were picked, grown, and purified using QIAprep Spin Miniprep Kit (Qiagen) and then sequenced by Sanger sequencing to confirm mutagenesis (Eurofins Genomics).

### Quantifying sleep behavior

30 dpf injected crispant fish and WT (uninjected) sibling controls were used for all behavioral experiments. Fish were maintained on a 10-14 light-dark cycle throughout development as well as for behavioral experiments. Fish were placed in 12-well tissue culture plates (Cellstar) 18-24 hours before behavioral recording to acclimate to the recording chamber. Fish were fed normal brine shrimp meals before the start of the recording, which began at ZT0 (zeitgeber time) and lasted 24 hours. Video recordings were processed in Ethovision XT (v13). Raw locomotor data was processed with custom-written scripts to quantify sleep duration and behavioral architecture such as locomotor distance, waking activity, sleep bout duration, and sleep bout number^38,74^. For each injected construct, three separate biological replicates were carried out to ensure the phenotype was consistently reproducible. Each biological replicate represented different clutches of fish, injected on different days. Both wildtype and crispant individuals were assessed from each clutch.

## Data accessibility

All sequence data were deposited in the short-read archive (SRA) under the accession PRJNA421208. Data underlying RNA FISH analysis can be found at http://www.stowers.org/research/publications/libpb-1485.

## Acknowledgements

We thank the University of Minnesota Genomics Center for their guidance and performing the cDNA library preparations and Illumina HiSeq 2500 sequencing. The Minnesota Supercomputing Institute (MSI) at the University of Minnesota provided resources that contributed to the research results reported within this paper. Funding was supported by NIH (1R01GM127872-01 to SEM and ACK) and NSF award IOS 165674 to ACK, and a US-Israel BSF award to ACK. KLM was supported by a Ruth Kirschstein National Research Service Award from NIH and a Stanford Center for Computational, Evolutionary and Human Genomics (CEHG) postdoctoral fellowship. CNP was supported by Grand Challenges in Biology Postdoctoral Program at the University of Minnesota College of Biological Sciences. Experiments were approved by the Institutional Animal Care and Use Committee at Florida Atlantic University (Protocols #A15-32 and #A18-38) and Stowers Institute for Medical Research (institutional authorization 2019-084).

## Author contributions

Conceptualization of experiments and analysis by SEM, ACK, NR, JK, KLM, JLP, JBJ; Methodology by SEM, KLM, JBJ, BAS, EF, DT, SES, BDS, JK, NR, AK, CNP; Formal analysis and investigation by KLM, JBJ, SEM, JLP, EF, DT, SES, BDS, JK; Writing – original draft by KLM, SEM; Writing – review and editing by JBJ, JLP, CNP, BAS, EF, DT, SES, BDS, JK, NR, ACK.

