## Supplemental Material for "Repeated evolution of circadian clock dysregulation in cavefish populations"

**Supplemental Material and Methods**

Supplemental Methods…………………………………………………………………...2-4
Supplemental References……………………………………..………………………6-8
Tables **S1-S11**……………………………………..………………………………9-20
Figures **S1-S14**……..……………………………………….……………………21-38

File S1

**Table S12**. Number of uniquely mapped reads per sample and sample annotations (population, timepoint)
**Table S13.** JTK_cycle results for each population
**Table S14.** Genes that are rhythmic in surface but arrhythmic in at least 1 cave (*p*>0.5)
**Table S15.** Curated list of genes with orthologs annotated as involved in circadian rhythm with their JTK_cycle *p*-values in cave and surface populations
**Table S16.** Pairwise $S_{DR}$ *p*-values between caves and surface
**Table S17.** Table S14. Phases called by JTK_cycle for genes across populations (includes genes without significant rhythmicity)
**Table S18.** Raw read counts for each sample
**Table S19.** Primer sequences
**Table S20.** Genetic differentiation at genes with changes in rhythmicity between surface and cave populations
**Table S21**. Pairwise comparisons of basal expression between surface and caves across timepoints with DESeq2
**Table S22**. Gene Ontology terms associated with cycling genes (FDR<0.05 in JTK_cycle) in each population

***Supplemental Methods***

***Sampling***
All samples from each population were derived from the same mating clutch and raised in 14:10 light-dark cycle. All fish were exactly 30 days post fertilization at the start of sampling. Fish were kept in total darkness for 24 hours prior to sampling and throughout the duration of the experiment. Fish were first sampled at 6am (Circadian Time 0) and then every four hours throughout until 2am (corresponding to CT 20). Prior to the experiment days, feedings were within a 2-3hr window to prevent fish from becoming food-entrained. Fish were fed *ad libitum* twice daily in the morning and evening (exactly at 8am and 8pm) during the experiment.

Six individuals were flash frozen separately for every time point and population and stored at -80°C until RNA was extracted.

***RNA extraction and library preparation***
For RNA isolation, all individuals were processed within a week of each other (between 1-19-2017 and 1-24-2017). Whole organisms (< 30 mg of tissue) were homogenized using Fisher brand pellet pestles and cordless motor (Fisher Scientific) in the lysate buffer RLT plus. Total RNA was extracted using the Qiagen RNAeasy Plus Mini Kit. Extraction batch was randomized across populations and treatments.

All cDNA libraries were constructed at the University of Minnesota Genomics Center on the same day in the same batch. In brief, a total of 400 ng of RNA was used to isolate mRNA via oligo-dT purification. dsDNA was constructed from the mRNA by random-primed reverse transcription and second-strand cDNA synthesis. Strand-specific cDNA libraries were then constructed using TruSeq Nano Stranded RNA kit (Illumina), following the manufacturer's protocol. Library quality was assessed using Agilent DNA 1000 assay on a Bioanalyzer. To minimize sequencing lane effects, barcoded libraries were pooled with treatment and population spread evenly between lanes. Samples were sequenced across multiple lanes of an Illumina HiSeq 2500 to produce 125-bp paired-end reads at University of Minnesota Genomics Center. All sequence data were deposited in the short-read archive (SRA).

***Read mapping and normalization***
Raw reads from each sample were assessed with Fastqc (v0.11.7), then cleaned of adapter contamination and low-quality bases with Trimmomatic (v0.33)[1]. Cleaned reads were mapped to the *Astyanax mexicanus* draft genome assembly version 1.02 (GCA_000372685.1) with STAR[2]. Counts for each gene were generated with stringtie (v1.3.3d)[3] based on the *Astyanax mexicanus* Ensembl v91 annotation. The Python script prepDE.py, bundled with stringtie, was used to generate a counts matrix for filtering, normalization, and subsequent analysis with DESeq2[4].

Genes with less than 100 total counts across all samples in the experiment were removed from the analysis. Counts were then normalized by sample library size and transformed with a variance stabilizing transformation with DESeq2. The variance stabilizing transformation makes the normalized counts values have approximately constant variance across mean expression values, which reduces bias due to highly expressed genes having greater variance than lowly expressed genes. Variance-stabilized counts were then decomposed for a principal component analysis (PCA) of all samples to identify broad patterns of differentiation among populations and timepoints. Principal components analysis was performed with the 'prcomp' package in the R computing environment.

***Promoter analysis of rhythmic genes***

Putative promoter sequences (1-kb upstream to 200bp downstream from the transcription start site (TSS)) were extracted based on the *A. mexicanus* genome assembly using samtools faidx. As the draft *A. mexicanus* genome assembly (v1.02) was built from an individual of Pachón cave descent, we used SNP and indel calls from population data[5] to create alternative reference sequences for Río Choy and cave populations by inserting variant calls into each reference sequence (see "Variant calling and population genetics measures" below). Variants where all genotyped individuals were homozygous for the alternative allele and at least 5 of 9 individuals were genotyped were inserted into the reference. LiftOver coordinates were created between alternative references to compare sequences between populations. When promoter regions, annotated based on the Pachón cave reference genome, were not available in an alternative population reference, the regions were not compared between surface-cave pairs. FIMO of the MEME suite[6] was then used identify motifs in each promoter. We used the default $p$-value cut-off of $< 0.0001$ of FIMO to identify significant motifs in each putative promoter sequence.

***Variant calling and population genetics measures***

Population genetic analyses were performed using samples from Herman *et al.*[5], using individuals from Río Choy (N=9), Pachón (N=9), Tinaja (N=10), Molino (N=9), as well as another surface population, Rascon (N=8). In brief, variant calling was performed using the Genome Analysis Toolkit v3.3.0 (GATK) based on GATK Best Practices[7,8]. Duplicate reads were marked using Picard's MarkDuplicates tool and filtered from downstream analyses resulting in a mean coverage of 9.28x. GATK's IndelRealigner (IR) and RealignerTargetCreator (RTC) were used to realign reads that may have been misaligned around insertions/deletions. HaplotypeCaller and then GenotypeGVCFs were then used to generate variant calls for all individuals. Low confidence variants were filtered using the VariantFiltration and SelectVariants tools.

To identify regions that were differentiated between cave and surface populations, we calculated multiple metrics based on the population variant calls. For population genomic metrics, we included the samples described above, as well as the Pachón reference genome[9], and required six or more individuals have data for a particular site. We excluded masked repetitive elements, indels, and the 10-bp surrounding indels. VCFtools v0.1.13[10] and custom scripts were used to calculate basic population genetic metrics ($\pi$, $F_{ST}$, and $d_{XY}$) in the coding region per gene. For $F_{ST}$ and $d_{XY}$ comparisons, we compared each cave population to two surface populations, Río Choy and Rascón.

Genes were considered $F_{ST}$ outliers if the average $F_{ST}$ of a gene was in the largest 5% of genes in the genome for a surface-cave comparisons (e.g., surface Río Choy vs. Pachón cave). As relative measures of divergence like $F_{ST}$ can be driven by low diversity due to low recombination or other genomic features[11], we also considered average $d_{XY}$. We considered genes as having a high $d_{XY}$ if average $d_{XY}$ was within the largest 20% of values for all genes in the genome for the focal comparison.

HapFLK v1.3[12] was used to estimate the HapFLK statistic for *Astyanax mexicanus* samples and two *Astyanax aeneus* samples sequenced in Herman *et al.*[5]. The Texas surface *Astyanax mexicanus* was excluded, as no population-level sampling was conducted for this population. HapFLK accounts for hierarchical population structure by building local ancestry and detecting changes in haplotype frequencies which exceed what is expected for selectively neutral evolution[12]. This method is thought to be robust to bottlenecks and migration and has performed well in comparison to other statistical tests for selection[13]. A population-specific tree was created for each scaffold with HapFLK prior to tests for selection. Reynolds distances and a kinship matrix were calculated with HapFLK. HapFLK was subsequently run on unphased data with 5 clusters (-K 5) and 20 EM runs to fit the LD model (-nfit = 20). *P*-values were estimated by fitting a standard normal distribution genome-wide. Genes with at least one $p$-value < 0.05 within the gene were subsequently compared to cave-surface $F_{ST}$ outliers to identify putative evidence for selection on the circadian transcriptome.

***Overlap with rhythmic transcripts in zebrafish***
To compare rhythmic transcripts in *A. mexicanus* with those of zebrafish, we compared genes with a JTK_cycle $p$-value < 0.05 to genes annotated as cycling in zebrafish in previous studies[14,15]. Genes annotated as significantly cycling under dark-dark and light-dark conditions in zebrafish larvae[14] and cycling in zebrafish liver under light-dark conditions[15] were downloaded and annotated to 1-to-1 orthologs of *A. mexicanus* genes for comparison.

***Annotating known circadian genes***
To create a list of known circadian genes, we downloaded 1-to-1 orthologs with zebrafish that were associated with the Go Ontology term "Circadian Rhythm." This list was also supplemented with genes from the literature[14]. The full list is found in File S1 (See tab "KnownCircadianPvalues"). Notably, three genes involved in the core feedback loop were not found to cycle significantly (at FDR<0.1) in any population: *clocka*, *clockb*, and *per2*. *Per2* is light activated and does not cycle robustly under darkness in zebrafish[14,16] (Figure S18). *Clocka/b* were lowly expressed in our dataset (average of 2.36 reads/sample and 12.8 reads/sample and for *clocka* and *clockb*, respectively), so we may lack power to infer rhythmicity at these genes.

***Identifying bases shifts between populations***
To identify base shifts (i.e., change in levels of expression but not necessarily a change in rhythm), we treated all timepoints as replicates and tested for differential expression with DESeq2 between pairs of populations (See File S1, tab "Baseshifts"). DESeq2 was used to normalize raw gene read counts, estimate dispersion factors for each gene, and then test for differential expression based on a negative binomial distribution with the default correction for multiple testing. To call differential expression, we retained genes at a cut-off of $p_{\mathrm{adj}} = 0.05$.

***Upregulation of DNA repair genes in cavefish populations***
Environmental light plays an crucial role in regulating DNA damage responses and cell cycle control in teleosts[17,18]. Previous work in the *A. mexicanus* system has found that cave populations show lower levels of DNA damage and increased expression of the DNA repair genes *CPD photolyas*e (*CPD phr*) and *damage-specific DNA binding protein 2* (*dd2*).[19] To explore this in

our dataset, we ask whether genes associated with the GO term for DNA repair (GO:0006281) showed higher basal expression in cave populations compared to the surface population in dark-dark conditions. We found that DNA-repair associated genes are upregulated in cave populations more than expected by chance (Fisher's exact tests, Table S10, all $p$-values < 0.0003 ).

***Comparison of basal expression of light-activated genes***
Previous work has shown that the light activate gene *per2* is upregulated in fin clips of cavefish from the Pachón and Chica caves[19]. The upregulation of this light-activated gene in cavefish has been suggested as evidence for constitutive upregulation of cavefish clock, similar to the condition of being under constant light[19]. To explore this question, we surveyed *A. mexicanus* orthologs of light-activated genes in zebrafish[14]. Light-activated genes are activated under the light phase in light-dark conditions, but maintain low levels of expression under dark-dark conditions[14]. Comparing basal levels of expression of this gene set across populations, we did not find that genes associated with light induction were more likely to be upregulated in cavefish compared to surface fish (hypergeometric tests, all $p$>0.84, Table S11). Nor do we find that light-activated genes in the core circadian feedback loop are consistently upregulated in cave populations. *Per2* showed higher expression in Tinaja ($q$=0.07) and Pachón ($q$=0.02), but was downregulated in Molino compared to the surface ($q$=0.004)(Figure S18). The light-activated gene *cry1aa*, which plays a key role in light entrainment[20], did not show differences in basal expression between cave and surface populations in our analysis (Tinaja, $q$=0.68; Molino, $q$=0.79; Pachón, $q$=0.82). Other cryptochromes which have been found to be light induced (*cry2a*, *cry3*)[21] also did not show differences in basal expression between cave and surface populations.

***Genetic differentiation at genes with rhythmic expression differences between cave and surface populations***
We asked whether genes in the circadian transcriptome showed genetic differentiation between surface and cave populations. Using population genomic data from the cave populations and two surface populations (Río Choy, Rascón) (see Methods), we found that 77 of the genes which showed reductions or loss of rhythmicity in at least one cave population were in the largest 5% of $F_{ST}$ values across all genes in the genome for at least one cave-surface comparison (File S1). Included in this set are genes known to be involved in circadian rhythm, such as *cry1ba*, *fkbp5,* and *pim3*[22]. Interestingly, the non-visual opsin gene exo-rhodopsin (*exorh*), which shows robust rhythmic expression in surface fish but is not significantly rhythmic in cavefish (Fig. S16), was in the top 5% of $F_{ST}$ values between Tinaja and both surface populations and Pachón and Río Choy. This gene also exhibited high absolute divergence ($d_{XY}$) for Tinaja to both surface populations, and showed high median $F_{ST}$ in comparisons between other cave-surface pairs (median $F_{ST}$, Río Choy-Molino: 0.46; Pachón-Rascon: 0.93). *Exorh* is the predominant opsin in the fish pineal gland and a candidate for mediating the effect of environmental light on pineal gene expression and melatonin synthesis[23–25].

Seven of the genes in this set of $F_{ST}$ outliers (*atp2b1b*, *gnb3b, dnajb9b, etv5b, cx52.7, opn3, ENSAMXG00000002780*) also showed signatures of selection based on the HapFLK statistic, which detects changes in haplotype frequencies which exceed what is expected under neutral evolution[12] (see Supplemental Methods). One of these, *dnajb9b,* a heat-shock protein homologous to human DNAJB9 which is under circadian control[26], also shows high absolute divergence between multiple cave and surface populations (File S1). Thus, we have some evidence that supports the notion genetic divergence observed in rhythmic genes may be shaped by selection.

***Animal care and use***

Procedures for all experiments performed at Florida Atlantic University were approved by the Institutional Animal Care and Use Committee at Florida Atlantic University (Protocols #A15-32 and #A18-38). Experiments performed at the Stowers Institute for Medical Research was approved by the Institutional Animal Care and Use Committee (IACUC) of the Stowers Institute for Medical Research. NR's institutional authorization for use of *A. mexicanus* in research is 2019-084.

**Table S1**. Numbers of rhythmic genes in each population

| | Number of rhythmic genes | |
|---|---|---|
| | FDR < 0.05 | FDR < 0.1 |
| Surface | 539 | 768 |
| Tinaja | 327 | 616 |
| Molino | 83 | 106 |
| Pachón | 88 | 193 |

**Table S2.** Number of genes with loss in rhythmicity ($P > 0.5$) in cave populations compared to rhythmic expression in surface (FDR < 0.1 and FDR < 0.05). 539 genes were rhythmic in the surface population at FDR < 0.05, and 768 genes were rhythmic in the surface population at FDR < 0.1.

| | 0.1 FDR | 0.05 FDR |
|---|---|---|
| Tinaja | 391 | 289 |
| Molino | 397 | 266 |
| Pachón | 393 | 252 |

**Table S3.** Known circadian regulators that are arrhythmic in one or more cave populations

| Gene | Arrhythmic |
|---:|---:|
| *Nfil3* | Pachón |
| *Aanat2* | Tinaja, Molino |
| *Cry1ba* | Tinaja, Pachón |
| *Cry1bb* | Tinaja, Pachón |
| *Arntl2* | Tinaja, Molino, Pachón |
| *Cry4* | Pachón |
| *Nptx2b* | Pachón, Molino |

**Table S4.** Phase (timing of peak expression) of core clock genes (primary and accessory loops) compared between zebrafish and *A. mexicanus* populations.

| | Zebrafish | Surface | Tinaja | Molino | Pachón |
|---|---|---|---|---|---|
| *Arntl1a* | 14.5 | 14 | 22 | 16 | 20 |
| *Arntl1b* | 13.8 | 14 | 20 | 16 | 16 |
| *Arntl2* | 17.4 | 18 | 8 | 22 | N/A[1] |
| *Per1a* | 2.3 | 2 | 6 | 6 | 6 |
| *Per1b* | 4.5 | 4 | 8 | 6 | 8 |
| *Rorca* | 12 | 12 | 14 | 12 | 10 |
| *Rorcb* | 13.3 | 12 | 14 | 16 | 14 |
| *Cry1ab* | 7.6 | 10 | N/A[1] | 8 | 0 |
| *Bhlhe40* | 5.3 | 4 | 8 | 8 | 8 |
| *Bhlhe41* | 1.3 | 0 | 6 | 2 | 4 |
| *Nfil3* | 16.2 | 12 | 2 | 14 | 4 |
| *Cry4* | 15.3 | 16 | 20 | 22 | 22 |
| *Cry1bb* | 15.4 | 18 | 2 | 18 | 0 |
| *Cry1ba* | 14.3 | 16 | N/A[1] | 20 | N/A[1] |
| *Nr1d1* | 0.8 | 0 | 6 | 2 | 4 |

[1]N/A indicates that amplitude is estimated at 0

**Table S5**. Phase shifts between surface and cave populations

| | Average shift | *P-value* |
|---|---|---|
| Surface-Pachón | 2.03 | $< 2.2 \times 10^{-16}$ |
| Surface-Tinaja | 1.3 | $< 2.2 \times 10^{-16}$ |
| Surface-Molino | 0.48 | $< 2.2 \times 10^{-16}$ |

**Table S6.** The number of significant circadian binding motifs identified in promoter proximal regions of rhythmic genes in the surface population.

| | Identified sequences | Number of genes |
|---|---|---|
| E-box(Arntl) | 426 | 296 |
| D-box(NFIL3) | 1240 | 602 |
| RORC | 776 | 775 |

**Table S7.** Arrhythmic genes in cave populations where motif sequences are also lost.

| Gene | Motif | Motif *p*-value in surface[1] | Matched sequence[1] | cave population in which motif is lost |
|---:|---:|---:|---:|---:|
| ENSAMXG00000015742 | D-box(NFIL3) | 8.81E-05 | TTTTGTAATCT | Molino |
| *yme1l1a* | D-box(NFIL3) | 3.73E-05 | TTATGTAAGTG | Molino |
| *gys1* | D-box(NFIL3) | 7.81E-05 | TTATATAATTT | Molino |
| *hdac1* | D-box(NFIL3) | 8.88E-05 | TTATATAATAA | Molino |
| *si:dkey-32e6.3* | D-box(NFIL3) | 3.9e-05;7.81e-05 | TTATATAACCT;TTATATAATTT | Molino |
| *edem3* | D-box(NFIL3) | 4.26E-05 | TTATATAATGT | Molino |
| *rab1ab* | D-box(NFIL3) | 3.91E-06 | TTATGTAATAT | Molino |
| *kdelr3* | D-box(NFIL3) | 4.01E-05 | TGATGTAACTT | Molino |
| *dhrs13a.2* | D-box(NFIL3) | 1.62e-05;3.02e-05 | TTACGTAACCA;TTACGTAAGCC | Molino |
| ENSAMXG00000027572 | D-box(NFIL3) | 4.8e-05;9.19e-05 | TTATATAATAT;TTATATAACTA | Molino |
| *ccdc187* | D-box(NFIL3) | 1.83e-05;2.92e-05 | TTACGTAACAG;TTACGTAAGTT | Molino |
| *si:ch211-191i18.4* | D-box(NFIL3) | 1.22e-05;8.2e-05 | TTATGTAATTT;TTACATAACAA | Molino |
| *si:ch211-157p22.10* | D-box(NFIL3) | 2.47E-05 | TTATGTAAGAA | Molino |
| *HOXC4* | D-box(NFIL3) | 9.86E-05 | ATATGTAATGC | Molino |
| *gata4* | D-box(NFIL3) | 1.83e-05;2.75e-05 | TTACGTAATAC;TTACGTAAGAC | Molino |
| *si:dkey-32e6.3* | D-box(NFIL3) | 3.9e-05;7.81e-05 | TTATATAACCT;TTATATAATTT | Tinaja |
| *ENSAMXG00000001997* | D-box(NFIL3) | 6.49e-06;1.83e-05 | TTACGTAATGT;TTACGTAATAC | Pachón |
| *ptges3a* | D-box(NFIL3) | 7.65E-05 | ATATGTAATAT | Pachón |
| *mapkapk2a* | D-box(NFIL3) | 8.97E-05 | TTTTGTAACAA | Pachon |
| *cry1bb* | D-box(NFIL3) | 4.80E-05 | TTATATAATAT | Pachón |
| *ENSAMXG00000019162* | D-box(NFIL3) | 2.58E-05 | TTATGTAATTA | Pachón |
| *si:dkey-17e16.10* | D-box(NFIL3) | 4.26E-05 | TTATATAATGT | Pachón |
| *mcm4* | E-box(Arntl) | 1.28E-05 | AGTCACGTGG | Molino |
| ENSAMXG00000008961 | E-box(Arntl) | 3.48E-05 | CCTCACGTGT | Molino |
| *pard6ga* | E-box(Arntl) | 1.28E-05 | AGTCACGTGG | Molino |
| ENSAMXG00000020281 | E-box(Arntl) | 5.87e-05;6.17e-05 | TAGCACGTGC;AAGCACGTGC | Molino |
| *gramd4b* | E-box(Arntl) | 4.63E-05 | CTTCACGTGC | Molino |
| *atad1b* | E-box(Arntl) | 3.95E-05 | TCTCACGTGT | Pachón |
| *tent5ba* | E-box(Arntl) | 2.41E-05 | AATCACGTGT | Pachón |
| *btr01* | RORC | 6.40E-05 | AAAACTGGGTTA | Molino |
| *ENSAMXG00000008655* | RORC | 8.49E-05 | GAAAATGGGTTA | Molino |
| *fdxacb1* | RORC | 5.85E-05 | TTAAACAGGTCA | Molino |
| *dpf3* | RORC | 1.87E-05 | AAAATTAGGGCA | Molino |
| *ENSAMXG00000009628* | RORC | 2.64E-05 | ATAAGCAGGTCA | Molino |
| *PAFAH1B2* | RORC | 8.72E-05 | GAAAGTATGTCA | Molino |
| *col8a1a* | RORC | 3.85E-06 | AAAAGTGGGTCA | Molino |
| *rpl7l1* | RORC | 3.76E-05 | AAAATGAGGTCA | Molino |

| | | | | |
|---|---|---|---|---|
| *zgc:92040* | RORC | 3.70E-05 | AATAGTAGGTCA | Molino |
| *SLC25A38* | RORC | 4.11E-05 | ATTATTAGGTCA | Molino |
| *si:dkey-275b16.2* | RORC | 9.88E-05 | ATAACTAGGTCT | Molino |
| *hyal2b* | RORC | 9.39E-05 | TATTATAGGTCA | Tinaja |

[1]When multiple motifs were identified in a proximal promoter, respective sequences and *p*-values are separated by a semi-colon.

**Table S8.** Average timing difference in peak expression of circadian feedback loop targets

| | Average difference in peak expression |
|---|---|
| | E-BOX (Arntl) |
| Molino-surface | 3.4 |
| Pachón-surface | 5.52 |
| Tinaja-surface | 7.24 |
| | RRE (Rorc) |
| Molino-surface | 4.4 |
| Pachón-surface | 5 |
| Tinaja-surface | 7 |
| | D-BOX (NFIL3) |
| Molino-surface | 4.11 |
| Pachón-surface | 5.23 |
| Tinaja-surface | 6.8 |

**Table S9**. Number of genes with a significant differential rhythmicity score

| | Number of differentially rhythmic genes (FDR<0.1) |
|---|---|
| surface vs. Molino | 148 |
| surface vs. Tinaja | 174 |
| surface vs. Pachón | 185 |

**Table S10.** Genes associated with GO term DNA-repair are upregulated in cave populations more than expected by chance. *P*-values based on Fisher's exact tests.

| | DNA-repair | Not DNA-repair | *p*-value |
|---|---|---|---|
| Upregulated in Molino | 28 | 3817 | |
| Upregulated in surface | 7 | 4204 | 0.0001 |
| Upregulated in Pachón | 70 | 3797 | |
| Upregulated in surface | 34 | 4038 | 0.0002 |
| Upregulated in Tinaja | 47 | 4653 | |
| Upregulated in surface | 10 | 5643 | < 0.00001 |

**Table S11.** *A. mexicanus* orthologs of genes that are light induced in zebrafish are not more often upregulated in cavefish compared to surface fish. *P*-values for each surface-cave comparison produced with a hypergeometric test.

| | Upregulated light-activated genes | Upregulated genes | Significance |
|---|---|---|---|
| Tinaja | 49 | 4700 | $p = 0.85$ |
| Surface | 70 | 5653 | |
| Pachón | 46 | 3867 | |
| Surface | 66 | 4072 | $p = 0.96$ |
| Surface | 63 | 4211 | |
| Molino | 37 | 3845 | $p = 0.99$ |

**Figure S1**. Raw reads per sample for Molino.

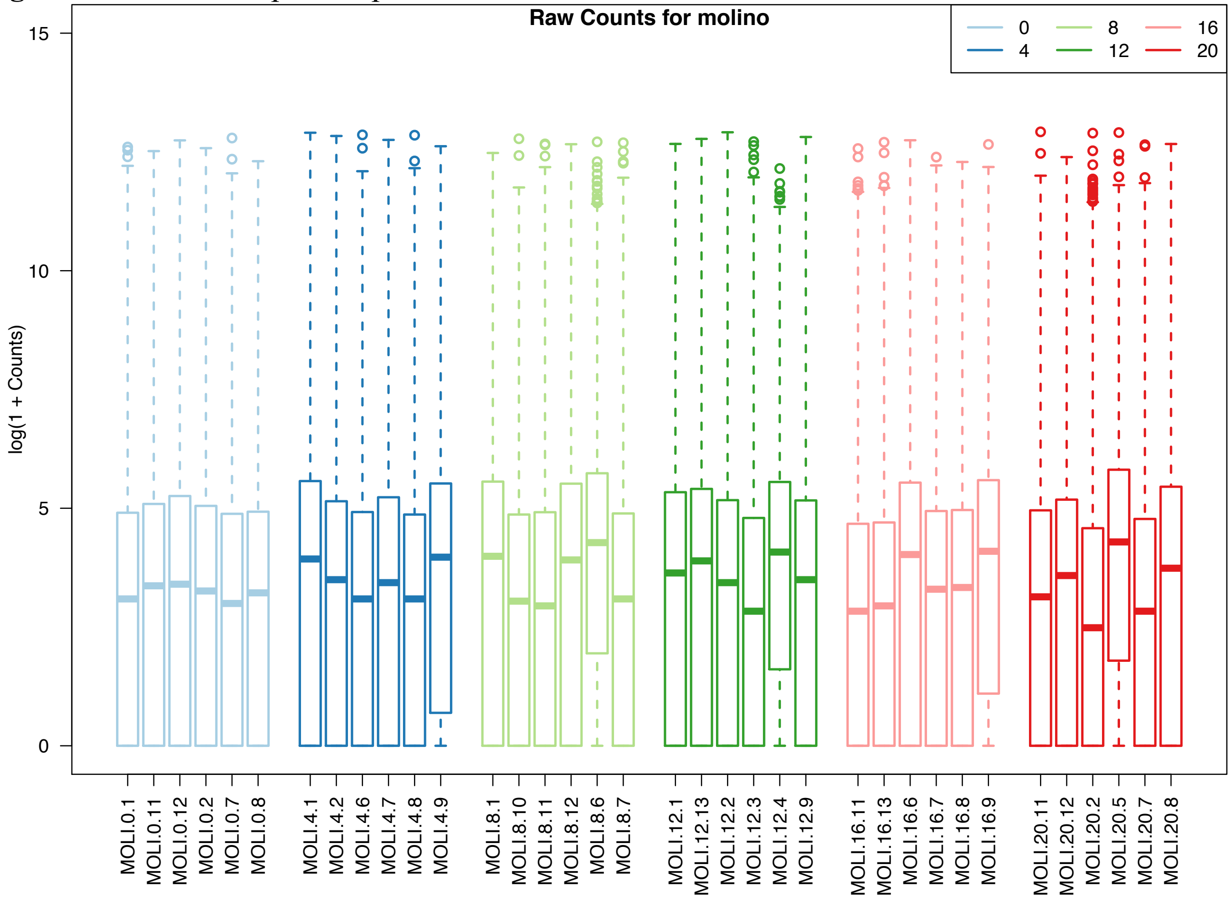

**Figure S2.** Raw reads per sample for Pachón.

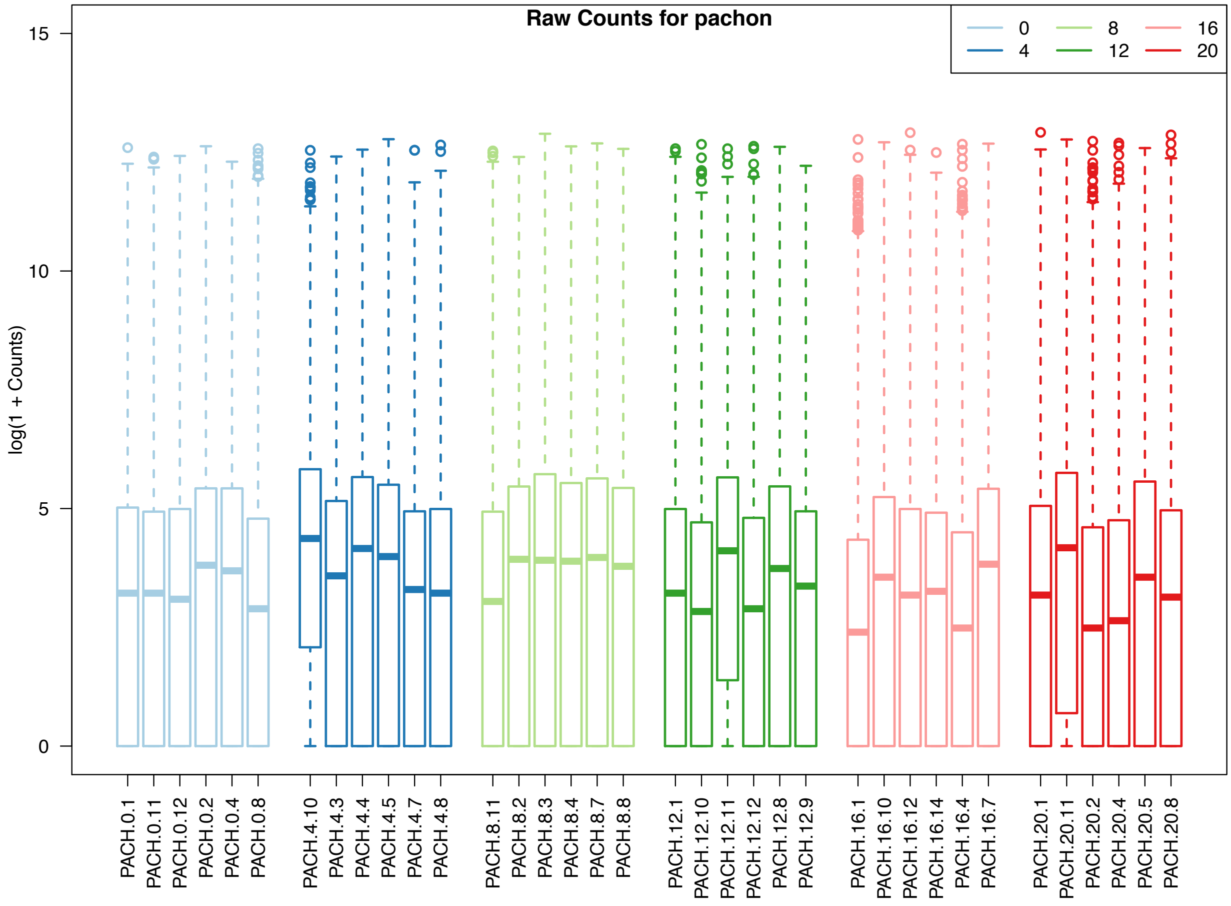

**Figure S3.** Raw reads per sample for surface fish.

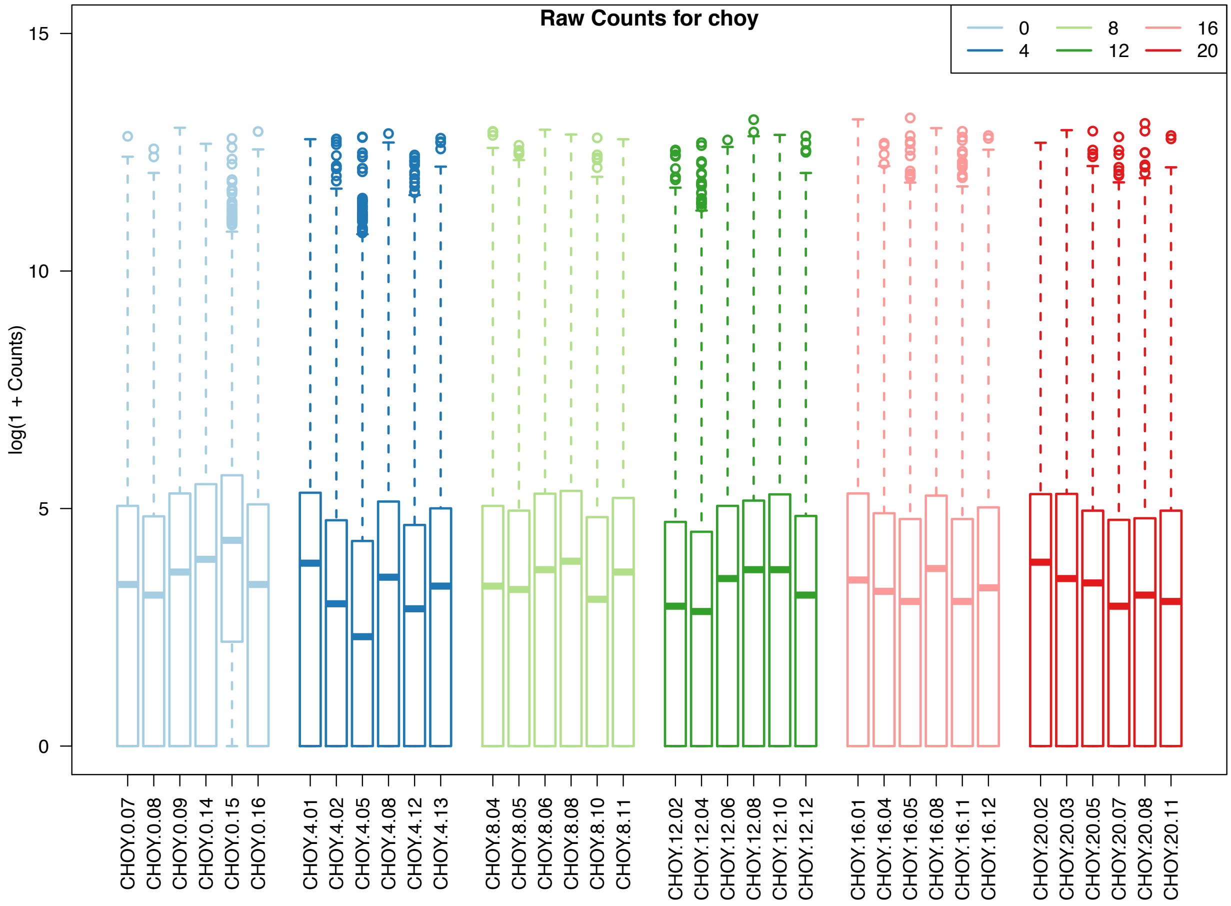

**Figure S4.** Raw reads per sample for Tinaja.

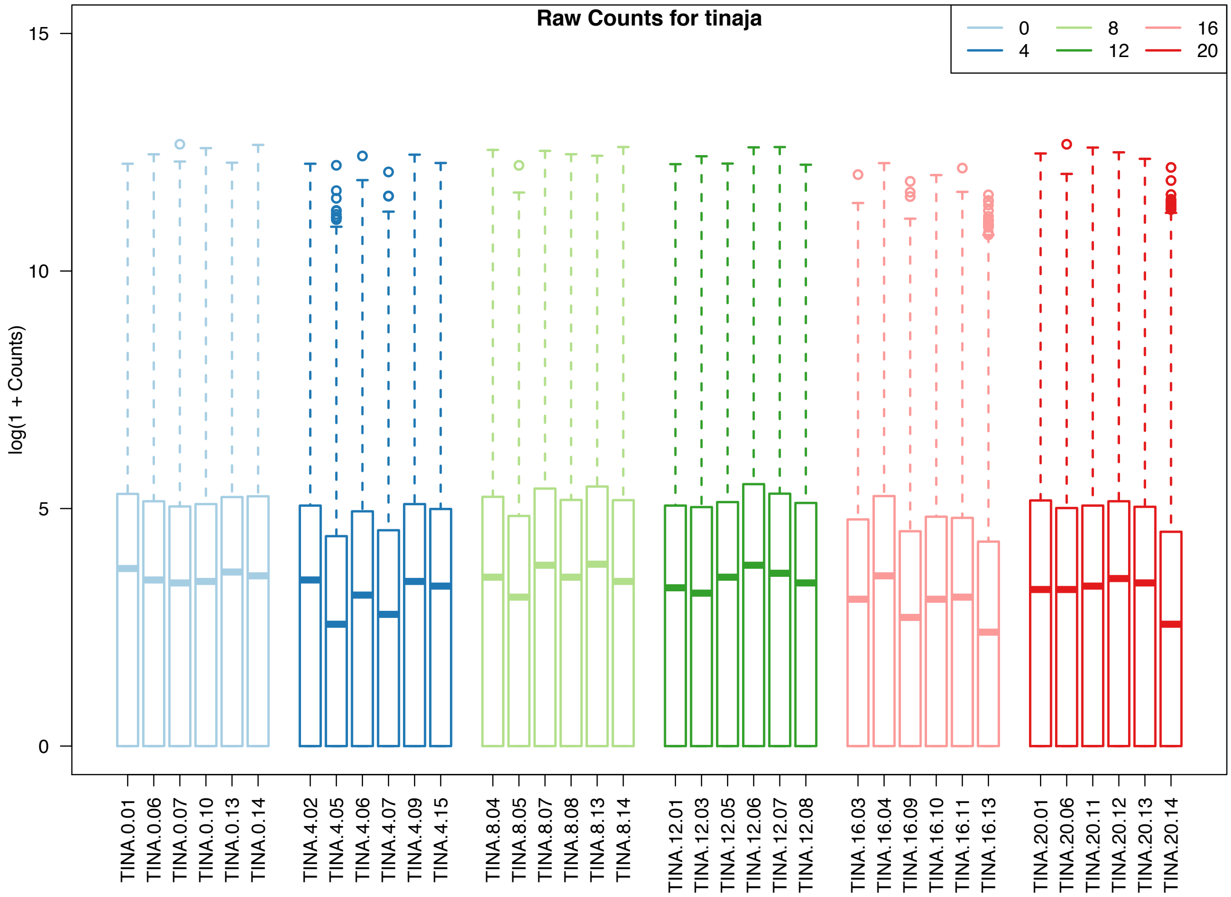

**Figure S5**. A. PC1 and PC2 (explaining 19.1% and 18% of variation, respectively) show that the primary axes of differentiation among samples is ecotype. B. PC3 (explaining 7.1% of variation) separates Molino from other populations.

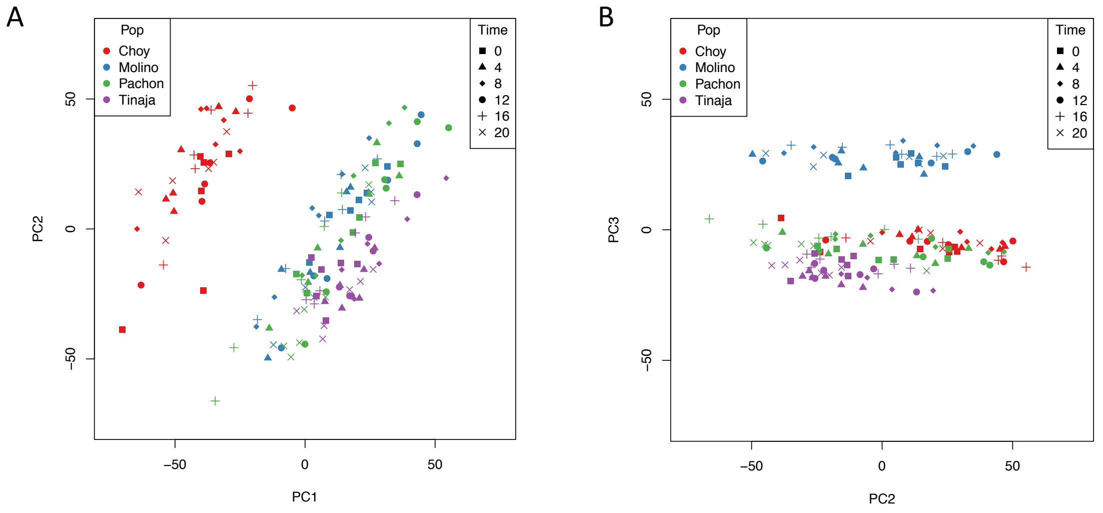

**Figure S6.** Cave populations show shifts in phase at *per1a/b* and *cry1a, w*ith gene expression peaking later in cave populations compared to the surface population.  Expression is represented as normalized read counts.

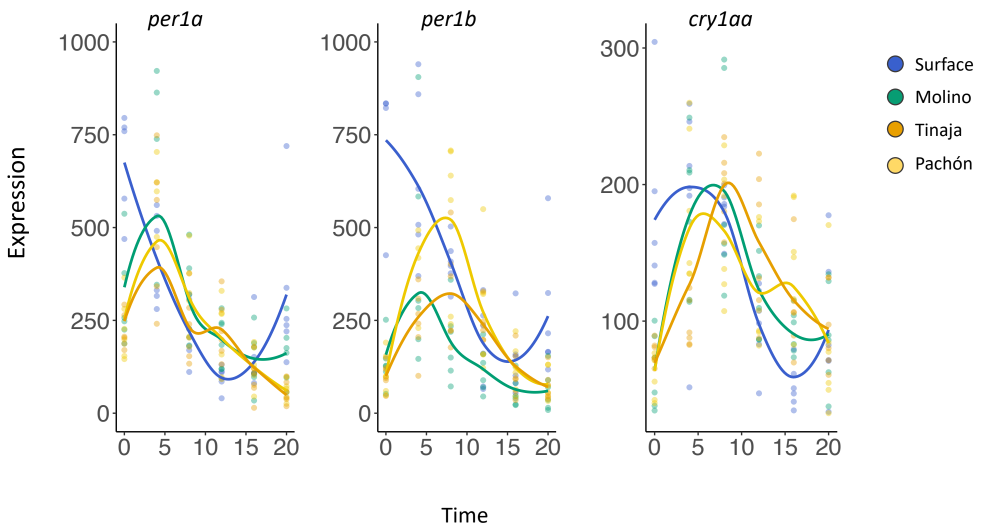

**Figure S7.** A-B. In the circles are the circadian phase distributions of predicted targets of RRE and E-BOX for surface fish. Grey bars represent the proportion of each motif seen in each phase. Highlighted in light grey are intervals of phase-specific enrichment for surface fish for each motif. Genes with the RRE motif (C) and EBOX (D) motifs show shifts in the timing of peak expression in cavefish populations.

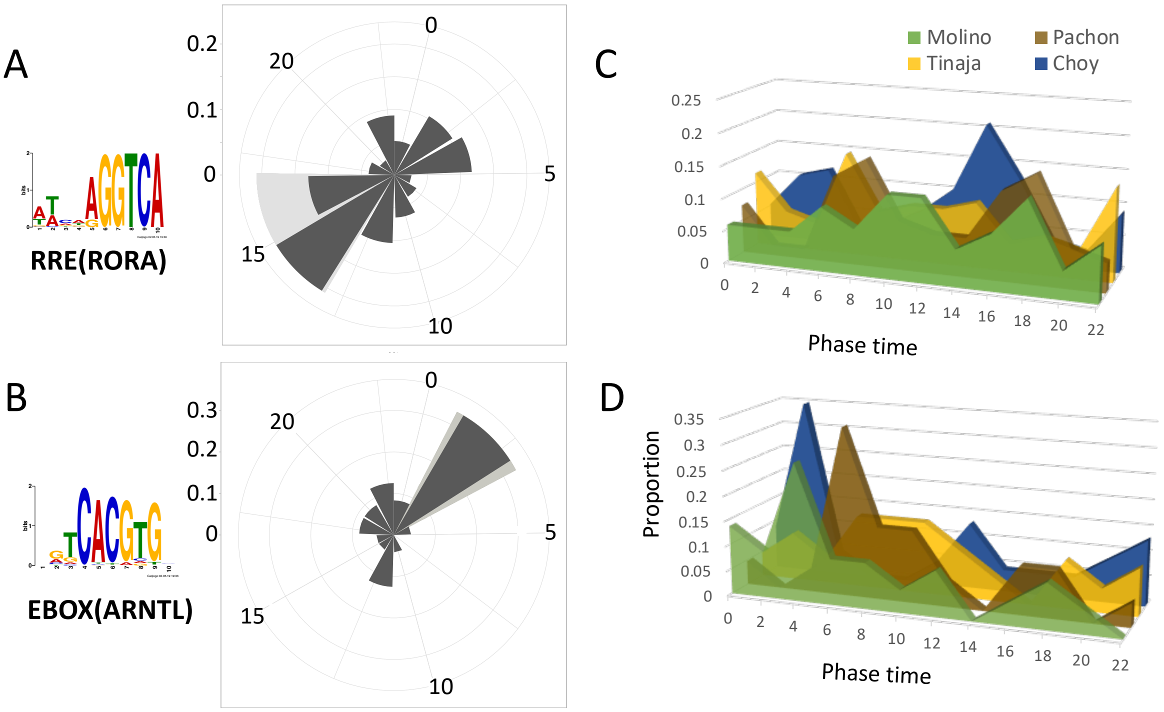

**Figure S8.** Region of interest used in all brain images (white box). Regions include the optic tectum (TeO) and periglomerular grey zone (PGZ) shown in relation to coronal section stained with DAPI. Scale bar is 50μM.

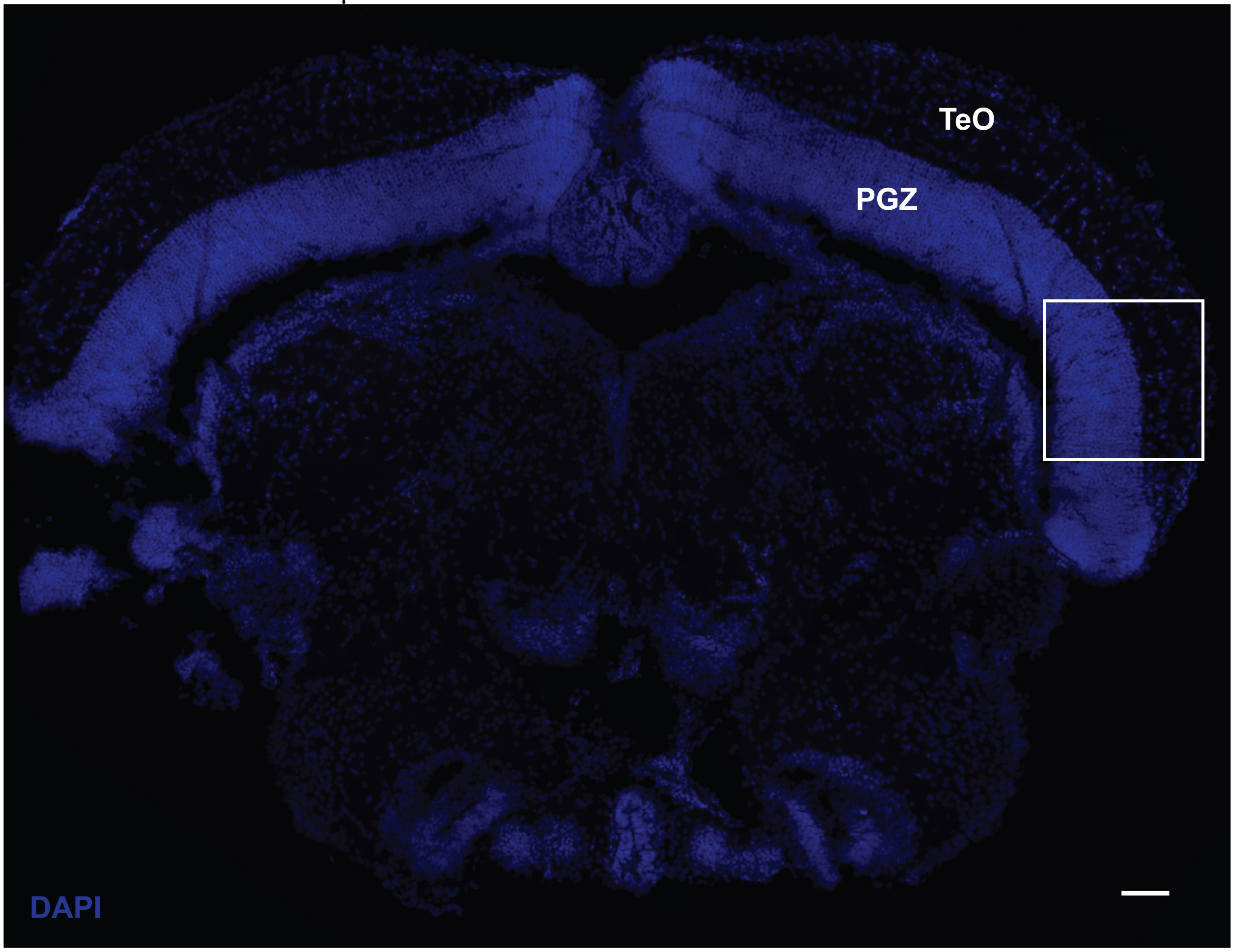

**Figure S9**. DAPI staining in brain ('B', top panels for each timepoint) and liver ('L', bottom panels for each timepoint) of surface fish and cavefish (Pachón, Tinaja, Molino) at CT0, CT8, and CT16. A. DAPI channel for sections included in Figure 4. B. DAPI channel for sections included in Figure 5.

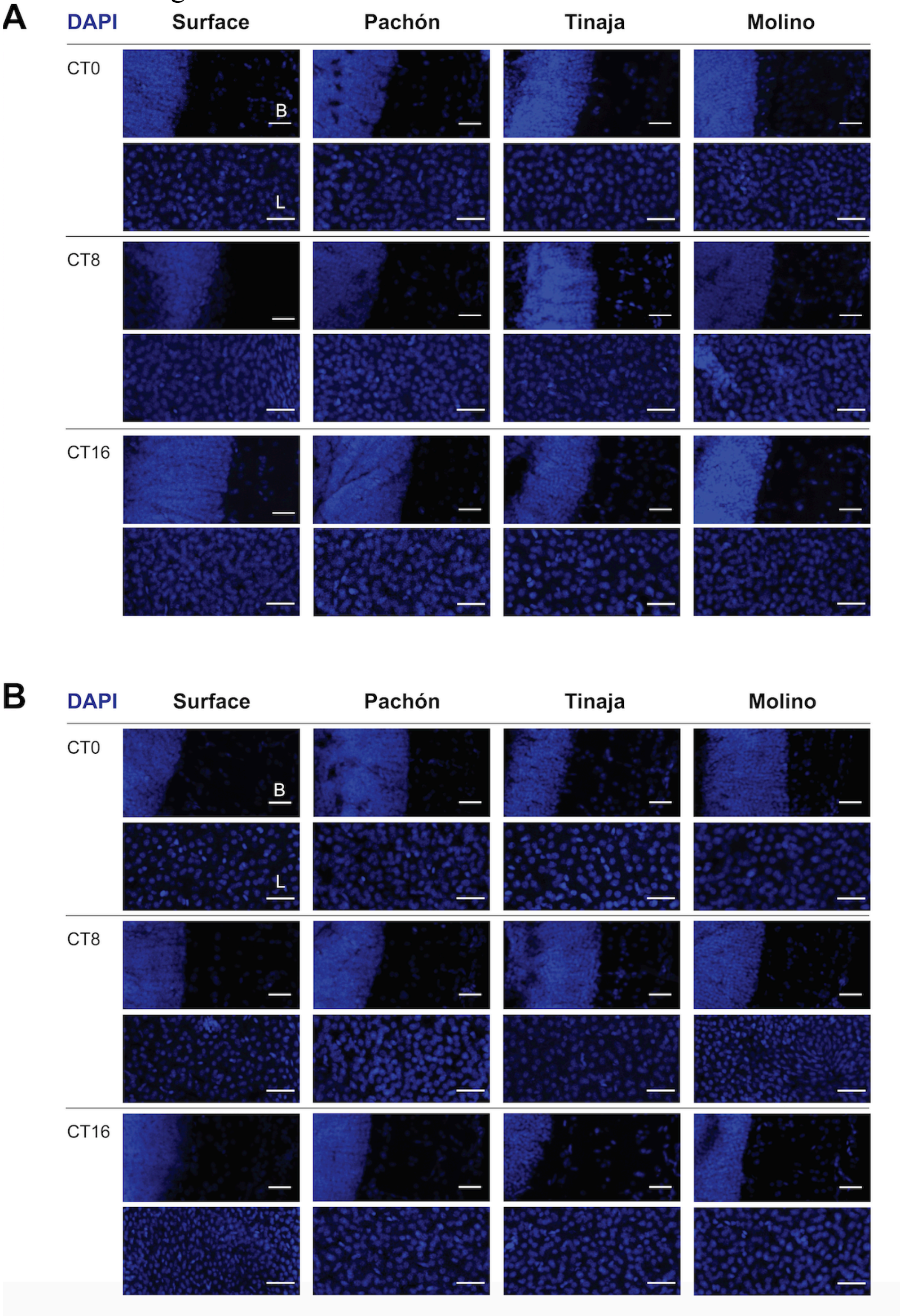

**Figure S10.** Quantification of RNA FISH. A-D. Expression of *per1a*, *arntl1a*, *rorca* and *rorcb* in brains (A, C) and livers (B,D) at CT0, CT8, and CT16 in surface fish and cavefish populations. RNAscope® probe channel intensity was normalized to DAPI channel intensity in identically sized, anatomically matched ROIs to provide an estimate of mRNA expression per cell (see Methods for full details of RNA FISH analysis). Biological replicates are shown as colored points on graph and represent a brain or liver sample collected from a single individual. Bars reflect mean and error bars show SEM of biological replicates. Statistics were calculated for each mRNA probe by comparing each cave population mean to control mean (Surface) within timepoints using ordinary 2-way ANOVA. Dunnett's test was used to correct for multiple comparisons across populations, timepoints. Adjusted p-values < 0.05 are reported with * using the following scheme: 0.0332 (*), 0.0021 (**), 0.0002 (***), and <0.0001 (****).

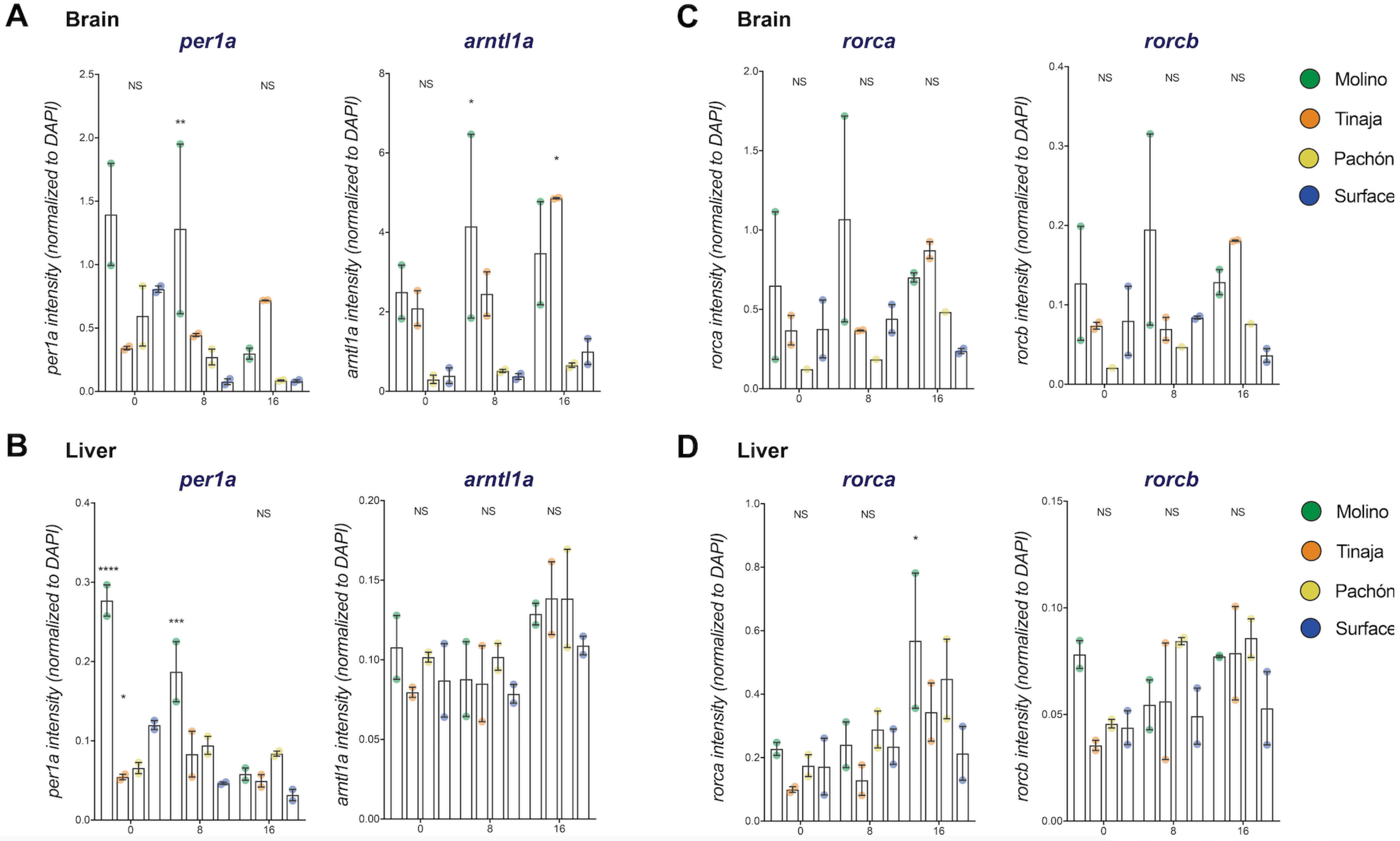

**Figure S11**. Expression patterns of *per1a* and *arntl1a* in Tinaja and Molino brain images adjusted to correct for oversaturation.

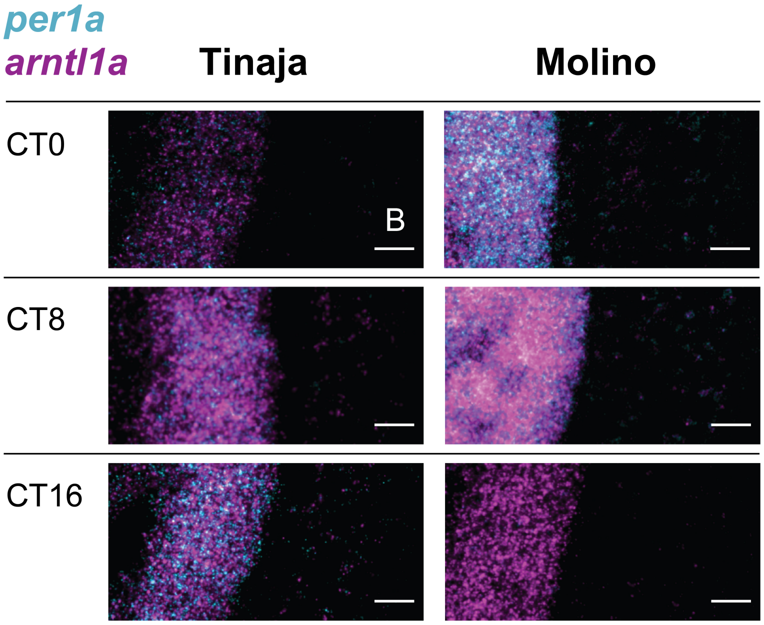

**Figure S12**. Temporal expression patterns of (A) *per1a* and (B) *arntl1a* in midbrain and liver tissue in *Astyanax mexicanus* populations. *In-situ* staining of *rorca* (A) and *rorcb* (B) using RNAscope® in midbrain ('B', top panels for each timepoint) and liver ('L', bottom panels for each timepoint) of Surface fish and cavefish (Pachón, Tinaja, Molino) at CT0, CT8, and CT16. Each time point is a single fish sample. Images are representative sections of two fish collected per time point, per population. Scale bar is 25μM.

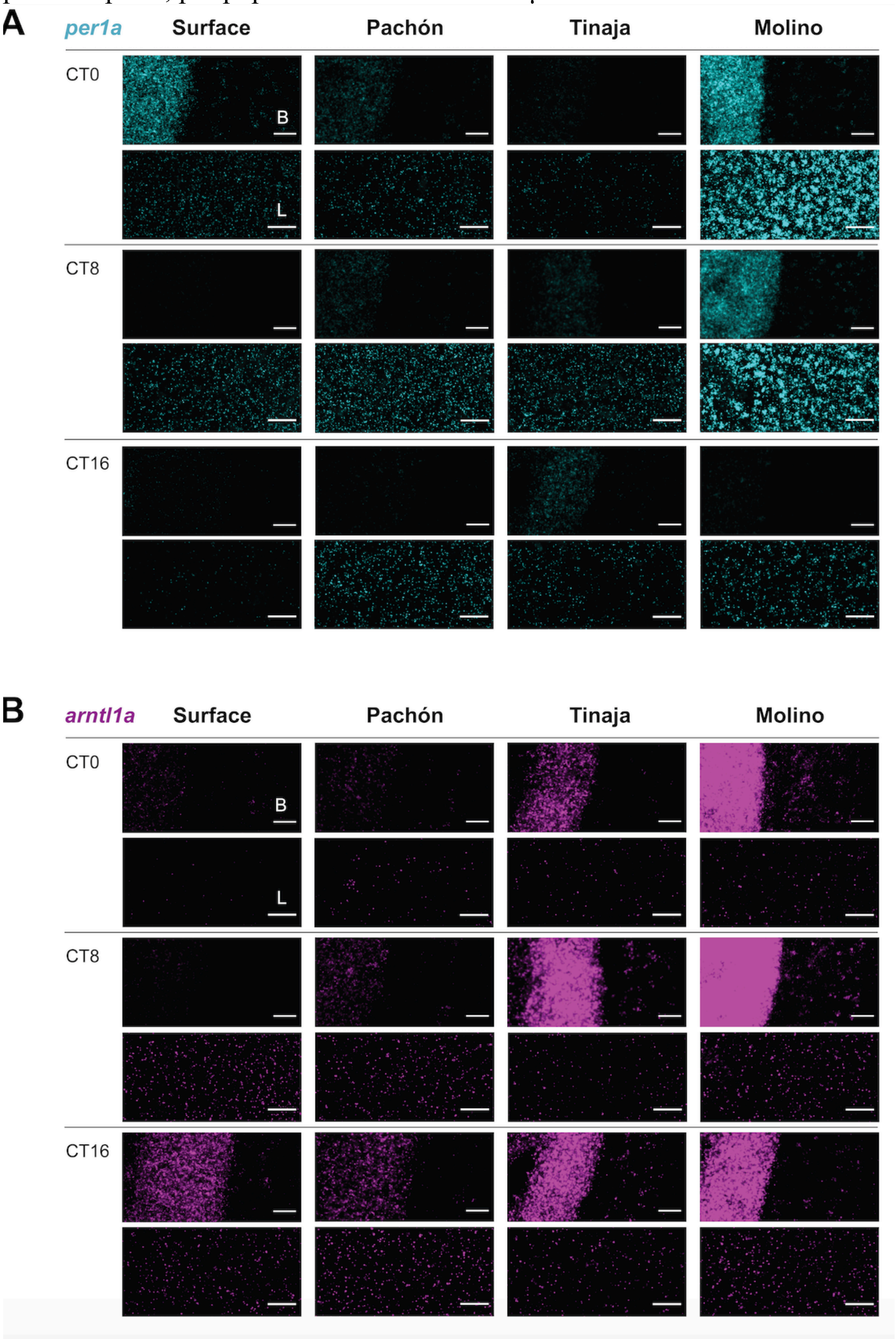

**Figure S13**. Temporal expression patterns of (A) *rorca* and (B) *rorcb* in midbrain and liver tissue in *Astyanax mexicanus* populations. *In-situ* staining of *rorca* (A) and *rorcb* (B) using RNAscope® in midbrain ('B', top panels for each timepoint) and liver ('L', bottom panels for each timepoint) of Surface fish and cavefish (Pachón, Tinaja, Molino) at CT0, CT8, and CT16. Each time point is a single fish sample. Images are representative sections of two fish collected per time point, per population. Scale bar is 25µM.

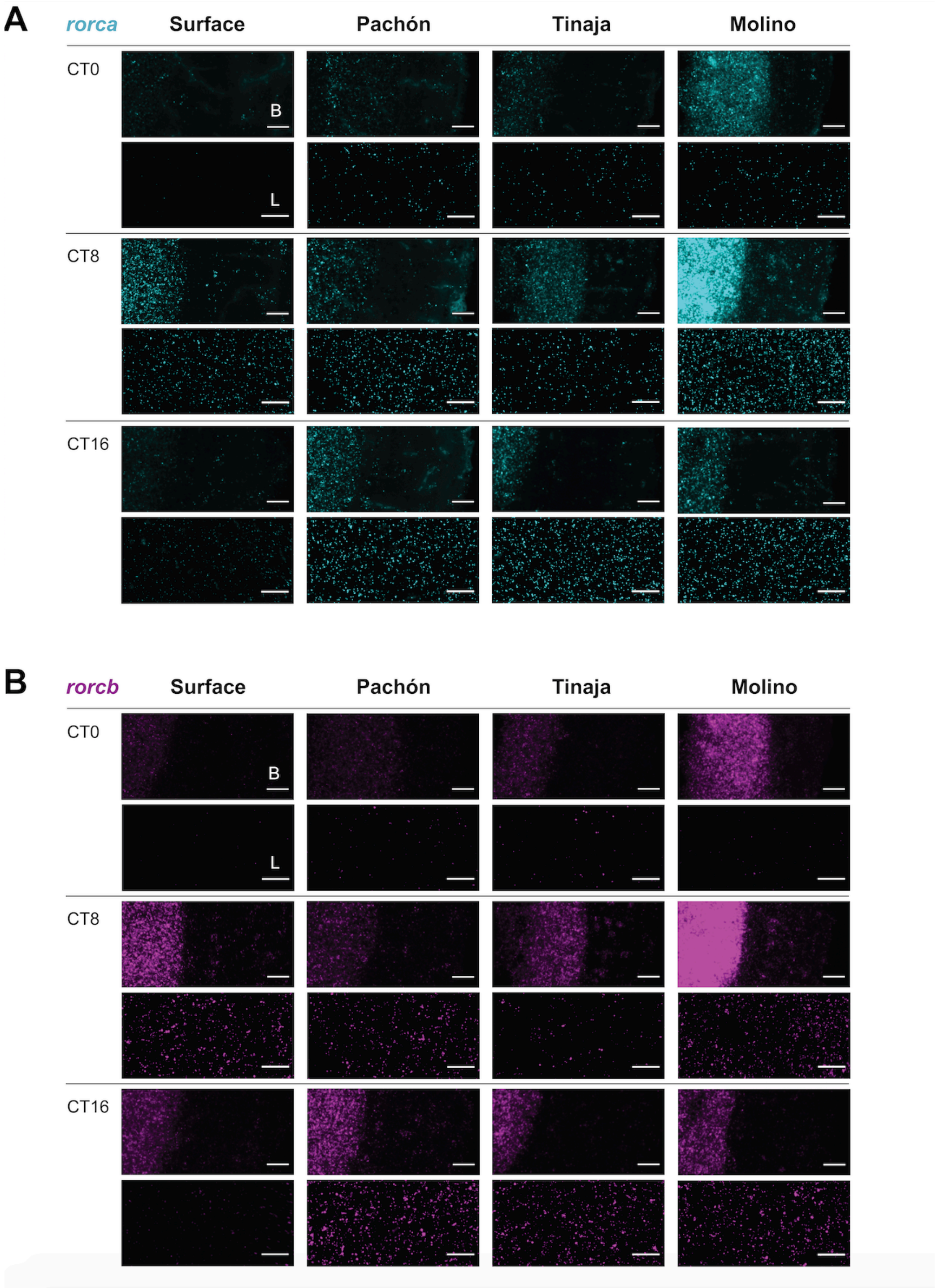

**Figure S14.** Analysis of mutagenesis in *aanat2* crispant $F_0$ fish. **A**. Genotyping gel of uninjected control and injected embryos. A portion of *aanat2* genomic region was amplified by PCR from DNA extracted from individual embryos. Labeled D is half of the PCR product that was digested with BbvI. Unlabeled is undigested PCR product. Indels can disrupt the restriction enzyme site, leading to undigested PCR product in injected embryos. **B**. Diagram of *aanat2* gene based on the surface fish reference genome (Ensembl v98). Boxes indicate exon and lines indicate introns. The empty boxes are 5' and 3' UTR and the closed boxes are coding sequence. A gRNA was designed targeting exon 1. The gRNA target site is in blue and the PAM sequence is in red. The underlined sequence is the BbvI restriction enzyme recognition sequence used for genotyping. The arrow indicates the predicted Cas9 cut site. Gene structure was generated using http://wormweb.org/exonintron and then modified. **C**. Sequence of wildtype surface fish and sequence of six clones from the restriction enzyme resistant band from *aanat2* injected individuals. The total number of base pairs less than the wildtype sequence is indicated to the right of each clone.

**A**

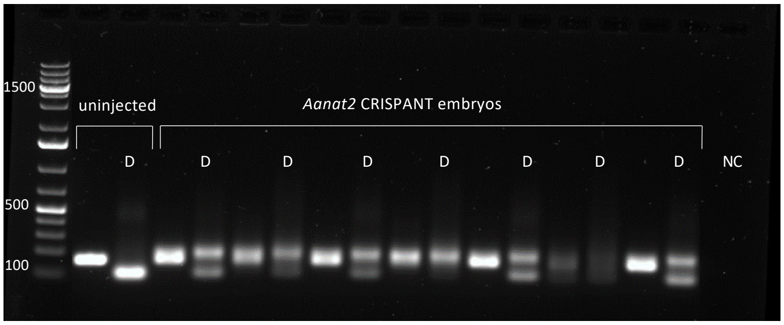


**B**

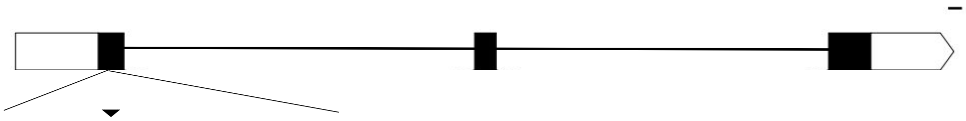

```
5'-GCCCCCTCCGGCAGCGGCGGCACACCCTGC-3'
3'-CGGGGGAGGCCGTCGCCGCCGTGTGGGACG-5'
```

**C**

```
CCAGCCCCCTCCGGCAGCGGCGGCACACCCTGCCCGCCAGCGAGTTCCGCAACCTCACGCCGCAGGACGCCATCAGCGTGTTCGAG   WT
CCAGCCC----------------------------GCCAGCGAGTTCCGCAACCTCACGCCGCAGGACGCCATCAGCGTGTTCGAG   -28
CCAGCCCCCTCCAGCGAGC--------------------------TCCGCAACCTCACGCCGCAGGACGCCATCAGCGTGTTCGAG   -23
CCAGCCCCCTCAGCG—GCGGCGGCACACCCTGCCCGCCAGCGAGTTCCGCAACCTCACGCCGCAGGACGCCATCAGCGTGTTCGAG   -1
CCAGCCCCCTCCAGCCCC-----CACACCCTGCCCGCCAGCGAGTTCCGCAACCTCACGCCGCAGGACGCCATCAGCGTGTTCGAG   -5
CCAG-----------------GGCACACCCTGCCCGCCAGCGAGTTCCGCAACCTCACGCCGCAGGACGCCATCAGCGTGTTCGAG   -17
CCAGCCCCC-----CAGCGGCGGCACACCCTGCCCGCCAGCGAGTTCCGCAACCTCACGCCGCAGGACGCCATCAGCGTGTTCGAG   -5
```

**Figure S15.** Analysis of mutagenesis in *rorca* crispant $F_0$ fish. **A**. Genotyping gel of uninjected control and injected embryos. A portion of *rorca* genomic region was amplified by PCR from DNA extracted from individual embryos. Labeled D is half of the PCR product that was digested with Cac8I. **B**. Diagram of *rorca* gene based on the Pachón Ensembl v93 genome. Boxes indicate exon and lines indicate introns. The empty boxes are UTR and the closed boxes are coding sequence. A gRNA was designed targeting exon 6. The gRNA target site is in blue and the PAM site is in red. The underlined sequence is the Cac8I restriction enzyme recognition sequence used for genotyping. The arrow indicates the predicted Cas9 cut site. Gene structure was generated using http://wormweb.org/exonintron and then modified. **C**. Sequence of wildtype surface fish and sequence of 3 clones from the restriction enzyme resistant band from *rorca* injected individuals. The total number of base pairs more or less than the wildtype sequence is indicated to the right of each clone.

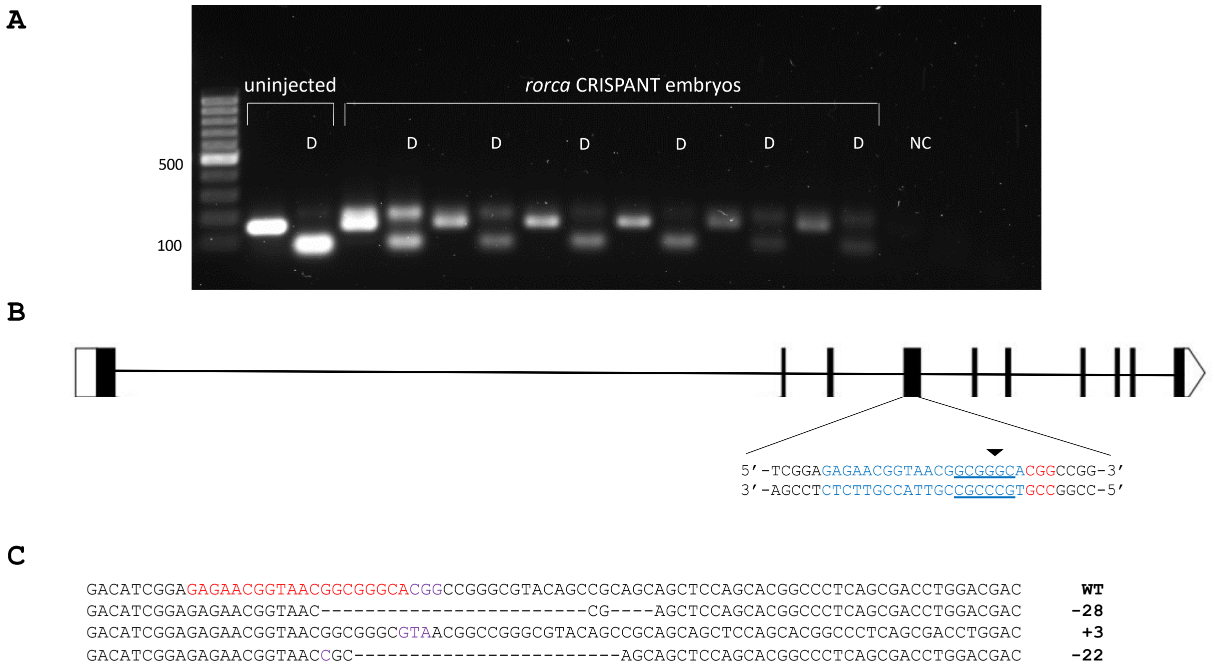

**Figure S16. A.** Expression of *per2*, a light-activated clock gene, over the course of the day in surface and cave populations (JTK_cycle *p*-values: surface, $p$=0.09, $q$=0.7; Molino, $p$=0.07, $q$=1; Pachón, $p$=0.003; $q$=0.16; Tinaja, $p$=0.016; $q$=0.22). Rhythmicity was not found to be different between populations (all comparisons, $S_{DR}$ $p$>0.68, $q$=1). **B.** Base level expression of *per2* between populations. *Per2* has lower base level expression in the surface than Pachón ($log_2$-fold change=0.82) and Tinaja ($log_2$-fold change=0.56), but higher expression than Molino ($log_2$-fold change=1.3).

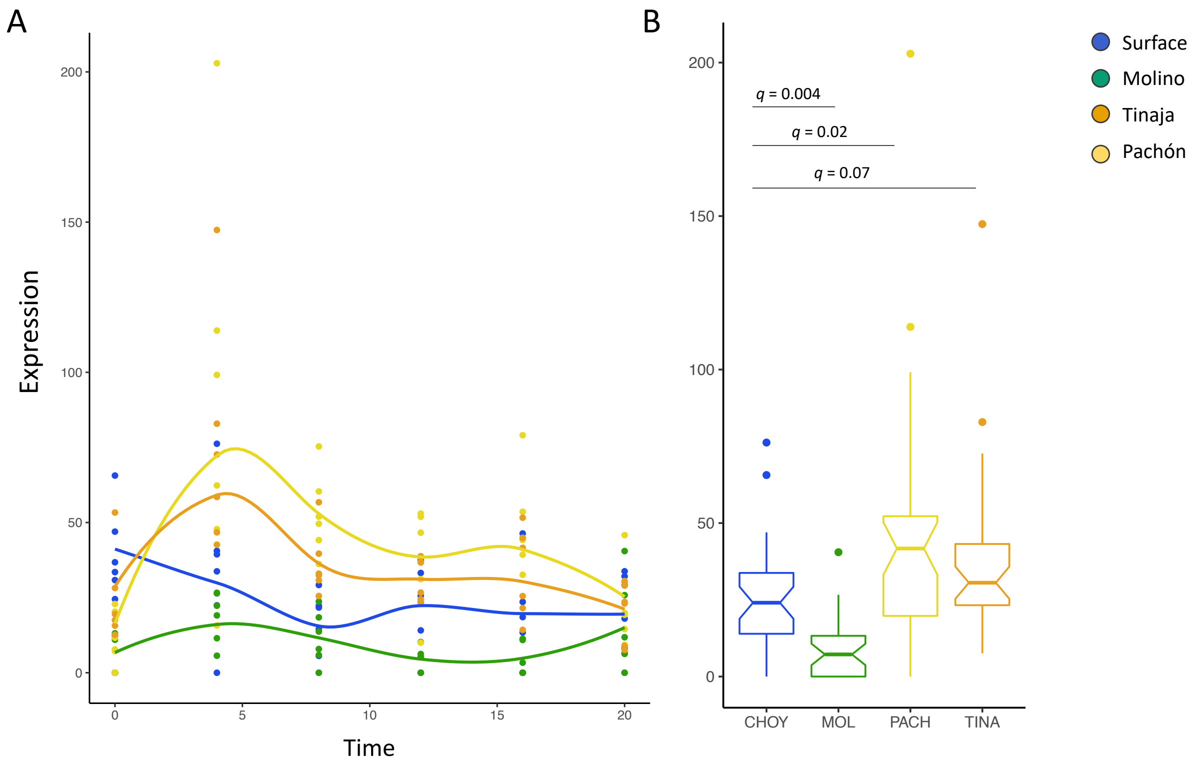

**Figure S17.** Core circadian genes and melatonin regulator *aanat2* show differentiated rhythmicity from surface fish in at least one cave population.

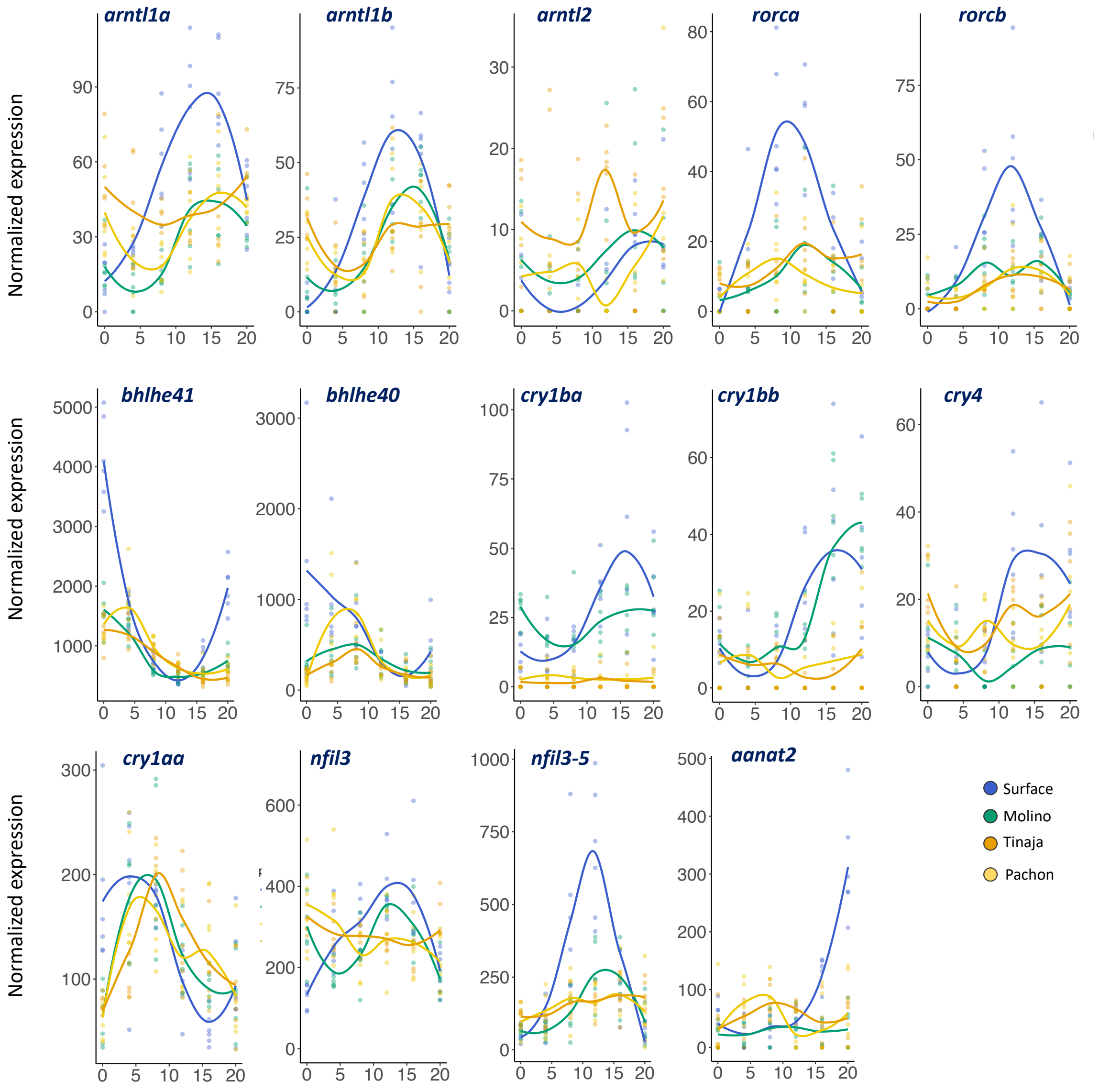

**Figure S18.** *Exo-rhodopsin* has robust rhythmic expression in the surface population ($p$=9.26 x $10^{-5}$, $q$ = 0.006), but is not strongly in rhythmic in cave populations (Pachón, $p$=0.04, $q$ = 0.32; Tinaja, $p$=1, $q$ = 1.0; Molino, $p$=0.37, $q$ = 1.0).

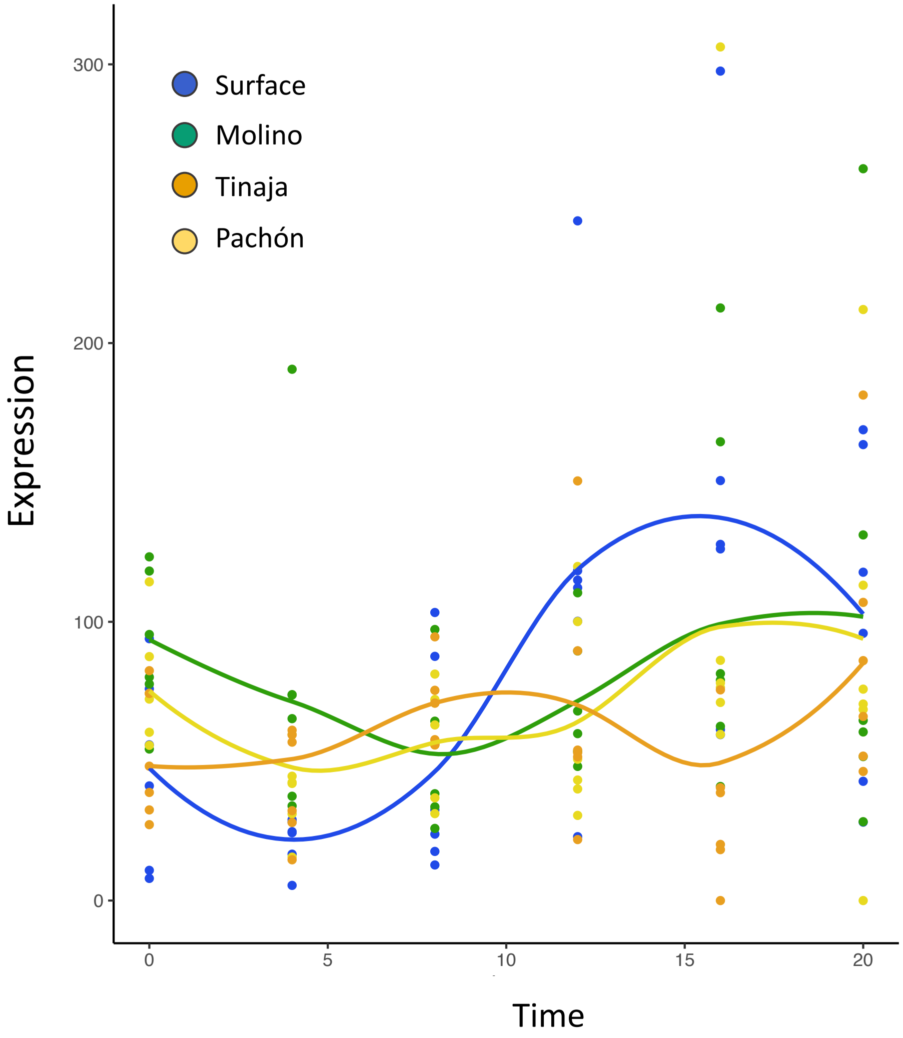
